# Genetically refactored *Agrobacterium*-mediated transformation

**DOI:** 10.1101/2023.10.13.561914

**Authors:** Mitchell G. Thompson, Liam D. Kirkpatrick, Gina M. Geiselman, Lucas M. Waldburger, Allison N. Pearson, Matthew Szarzanowicz, Khanh M. Vuu, Kasey Markel, Niklas F. C. Hummel, Dennis D. Suazo, Claudine Tahmin, Ruoming Cui, Shuying Liu, Jasmine Cevallos, Hamreet Pannu, Di Liu, Jennifer W. Gin, Yan Chen, Christopher J. Petzold, John M. Gladden, Jay D. Keasling, Jeff H. Chang, Alexandra J. Weisberg, Patrick M. Shih

## Abstract

Members of *Agrobacterium* are costly plant pathogens while also essential tools for plant transformation. Though *Agrobacterium*-mediated transformation (AMT) has been heavily studied, its polygenic nature and its complex transcriptional regulation make identifying the genetic basis of transformational efficiency difficult through traditional genetic and bioinformatic approaches. Here we use a bottom-up synthetic approach to systematically refactor the tumor-inducing plasmid, wherein the majority of AMT machine components are encoded, into a minimal set of genes capable of plant and fungal transformation that is both controllable and orthogonal to its environment. We demonstrate that engineered vectors can be transferred to new heterologous bacteria, enabling them to transform plants. Our reductionist approach demonstrates how bottom-up engineering can be used to dissect and elucidate the genetic underpinnings of complex biological traits, and may lead to the development of strains of bacteria more capable of transforming recalcitrant plant species of societal importance.

## Introduction

The genetic basis of pathogenesis is challenging to study due to its highly polygenic nature as well as it being influenced by both host and environmental factors ^1^. While advances in comparative and functional genomics have generated myriad hypotheses on how virulence and adaptations to specific hosts evolve ^2,3^, it is still challenging to isolate and validate specific genetic features that determine these traits ^4^. In an ideal system, one would be able to systematically evaluate and build a holistic understanding of how each gene contributes and influences virulence. However, epistatic effects often complicate the conclusions drawn from traditional top-down approaches that rely solely on knockouts and complementation ^5^.

As an alternative bottom-up approach, synthetic biology enables the introduction of synthetic regulatory control on a defined set of genetic elements. This is crucial for two major reasons. First, the development of minimal, controllable systems allows for specific hypotheses to be tested to better understand how evolution has solved a myriad of problems. Second, this knowledge gained allows for subsequent data-guided engineering to optimize and leverage the system for biotechnological purposes. Such a strategy has been widely implemented in reconstituting relatively linear metabolic pathways ^6,7^, but apart from a few notable exceptions, it has rarely been applied to more complex biological phenomena because of the numerous and tremendous intrinsic challenges associated with building up complex biological traits in a reductionist manner ^8,9^. To perform “genetic refactoring” one must identify the genes necessary and sufficient for a given biological process ^5^, as well as have the appropriate genetic tools applicable to the organism of study^10^. Given these significant hurdles, many initial designs from genetic refactoring often perform poorly compared to their native system, but nonetheless offer unique insights into the underlying complexities facets of biological traits ^9,11^.

A problem unique to studying the genetic bases of pathogenesis is that any synthetic regulatory elements utilized must also be robust *in situ*, *i.e.*, in the context of the various environments that the pathogen faces during infection, where very few genetic toolkits have been rigorously validated. Despite these challenges, work with both plant-and mammalian-associated bacteria has demonstrated that synthetic genetic constructs can be introduced to promote non-native interactions ^12,13^, indicating the feasibility of a complete synthetic refactoring of pathogenesis. Nonetheless, genetically recapitulating complex biological phenomena within a host-associated environment has largely remained out of reach.

Plant pathogenic members of *Agrobacterium* and *Rhizobium* (hereafter collectively referred to as *Agrobacterium tumefaciens*) are capable of causing crown gall or hairy root diseases and have been extensively studied due to their unique mechanisms of virulence. Virulence involves genetic transformations of eukaryotic hosts, which has been leveraged for many critical biotechnological uses, *e.g.*, plant transgenesis^14^. Central to virulence is an oncogeneic tumor-inducing plasmid (pTi) that carries a “Transfer DNA” (T-DNA) and *vir* genes. The hallmark of *A. tumefaciens* virulence is the transfer of a protein-conjugated, single-stranded DNA molecule into host cells and integration of the DNA into the genome. When genes from this T-DNA are expressed in the genetically modified plant cell, the gene products synthesize phytohormones that result in the formation of a tumor. The infecting bacterial population is hypothesized to gain a fitness advantage in the tumor because of access to novel nutrients, which are also encoded for on the T-DNA ^15^. When scientists domesticated virulence by swapping the tumorigenic genes within the T-DNA region with genes of interest, a new era of plant genetics was ushered in. Today the T-DNA borders and genetic payloads to be delivered are most often housed on a smaller plasmid referred to as a binary vector, enabling easy genetic manipulation through *Agrobacterium-*mediated transformation (AMT) of multitudes of plant and fungal species ^16–18^. However, many agriculturally important crops still remain difficult to transform^19^. Thus, there remains a tremendous imperative to develop novel strains of *Agrobacterium* that will enable scientists to expand the genetic potential of plants.

Our basic understanding of AMT and nearly all of the engineered agrobacterial strains used for AMT are derived from a limited number of *A. tumefaciens* strains and pTi variants ^20^. Yet, it has long been recognized that interactions among strains, Ti plasmids, and host species influence the efficiency of AMT^21^. By mining this natural diversity, strains with improved plant transformation properties for different plant species have previously been developed ^22,23^. More recently, groups have developed strains that contain additional *vir* alleles, harbored either on the binary vector (superbinary vectors) or on an additional stand-alone plasmid (ternary vectors)^24–26^. These strains demonstrate that altering the regulation of *vir* genes can enhance transformation of otherwise recalcitrant plants ^24–26^. Precisely how these tripartite interactions influence transformation efficiency remains largely unknown. The high number of possible genetic interactions required for AMT complicates research efforts at improving transformation by *A. tumefaciens*. Complicating studies is that oncogenic plasmids vary in the composition and sequence of *vir* genes, the regulation of these genes^27,28^, and that chromosomal genes implicated in virulence vary in sequence across agrobacterial strains ^29–31^. Furthermore, the impact of changes in expression level between different *vir genes* is somewhat masked by a master regulator, VirA/G, which controls the expression of all known *vir* genes ^30–32^. This epistatic regulatory schema makes it difficult to evaluate whether differences in virulence are a consequence of the presence of a specific *vir* gene or its strength of expression ^33,34^. Thus, to fully capture the impact of these many individual genetic variables involved in AMT, a bottom-up synthetic genetic approach is required to precisely control genetic interactions and systematically evaluate the contribution of each gene to transformation. However, due to the sheer size of combinatorial genetic space within pTi that can be explored and the technical challenges associated with refactoring complex biological phenomena *in planta*, no such effort has been reported.

Despite the many technical challenges associated with engineering synthetically encoded AMT, a deeper understanding of this complex process may elucidate molecular constraints to the transformation of plants. Here we overcome these challenges by 1) developing a set of genetic tools that allow for the reliable control of agrobacterial gene expression within the plant environment, in order to 2) quantitatively characterize the genetic determinants underlying AMT, to ultimately 3) design synthetic vectors, divorced from native regulation, capable of plant transformation. This work represents a critical first step in better understanding AMT as we lay the framework for understanding highly specific genotype-to-phenotype connections in a complex host-microbe interaction.

## Results and Discussion

### Developing a genetic toolkit to control bacterial gene expression in planta

A recurring challenge in synthetic biology has been translating genetic circuits developed *in vitro* into more heterogeneous environments *in situ*. Environmental changes can have dramatic impact on genetically engineered organisms, as demonstrated in scaleups to large fermentative tanks or living medicines in patients ^35,36^. Many *in vitro* synthetic biology designs take advantage of small-molecule inducible promoters, which offer a range of expression options from a single design, compared to static expression levels from a single constitutive promoter. However, dynamic environments such as plant tissue may interfere with inducible promoter systems by making signaling molecules biologically unavailable through degradation or sequestration, thus dramatically limiting their potential usefulness. Recent work by multiple groups <u>have</u> characterized inducible promoters in *Agrobacterium*, though not all were evaluated while the bacteria was *in planta*; moreover, there has been a dearth of well characterized constitutive promoters in *Agrobacterium*^37–39^.

To better understand how to control bacterial gene expression within plants, we evaluated the activity of 16 synthetic constitutive, and 4 inducible promoters in bacterial cells grown in rich media ^40^, as well as infiltrated into the leaf tissue of *Nicotiana benthamiana* and *Arabidopsis thaliana*. In both plants, bacterial constitutive promoter activity correlated highly between leaf tissues, and between observed *in vitro* activity **(Figure 1A, Figure S1)**. We then choose five constitutive promoters with a range of expression strengths (P_J23114_, P_J23117_, P_J23101_, P_J23100_, and P_J23111_) to complement a *virE12* deletion mutation in the common *Agrobacterium fabrum* (formerly *A. tumefaciens*) laboratory strain GV3101, which is derived from *A. fabrum* C58. Previous work using inducible promoters showed that transformation of tobacco was highly sensitive to *virE12* expression ^39^. Using transiently expressed GFP from a medium strength plant promoter in *N. benthamiana* as a measure of transformation ^41^, we observed that all promoters stronger than the weakest, P_J23114_, were able to complement leaf transformation back to wild-type levels (**Figure 1B**). Proteomics analysis of the Δ*virE12* complementation strains confirmed that expression of VirE12 correlated with RFP expression from the same promoters (**Figure 1C**). These data reveal that relatively weak constitutive promoters may be sufficient to reconstitute *vir* gene expression from pBBR1 origin plasmids.

**Figure 1:**
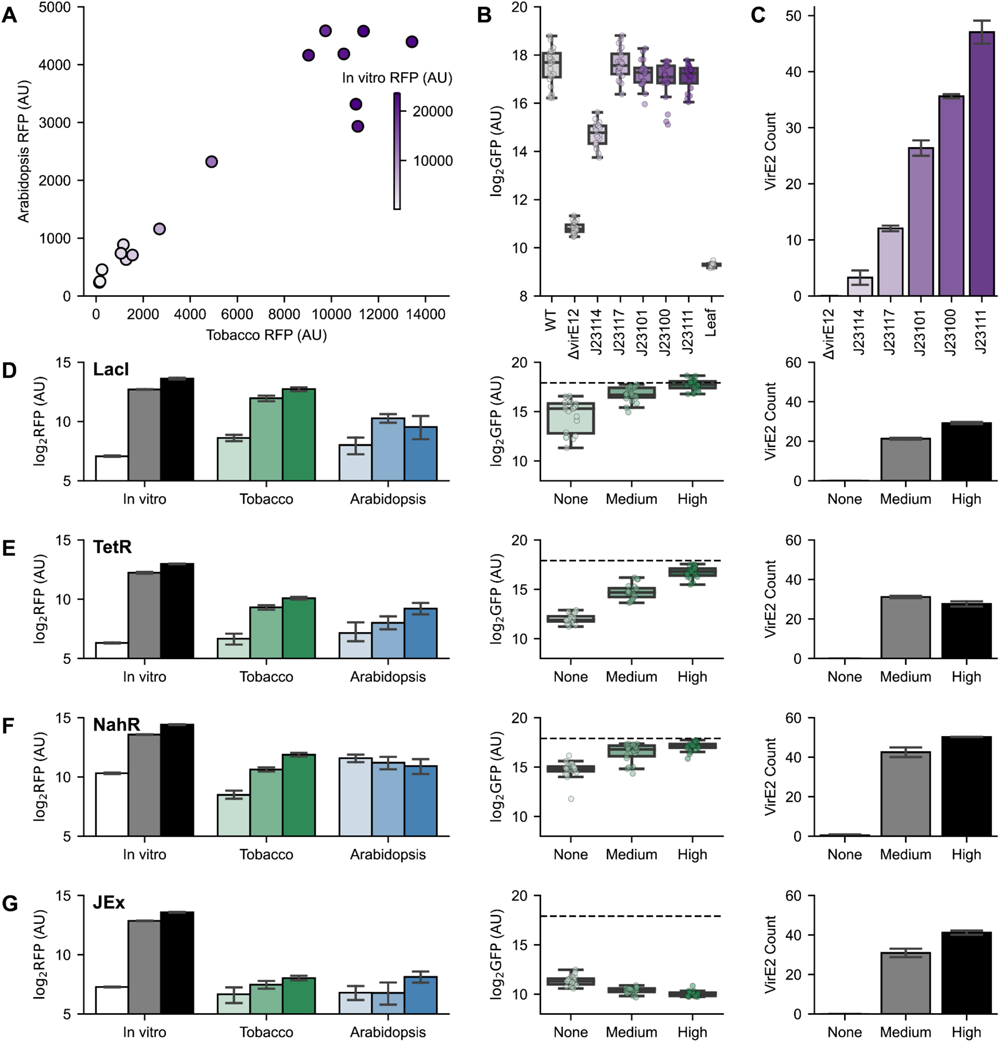
Characterizing a Synthetic Biology Toolkit *in planta*: A) Activity of constitutive promoters driving RFP from pGingerBK plasmid backbone. The x-and y-axes show RFP production from *A. fabrum* C58C1 3 days after infiltration in tobacco or *Arabidopsis* respectively (n=12). The color palette displays the activity of the same promoter *in vitro* (n=8). B) Transient expression of GFP from agroinfiltrated tobacco leaves (AU) log2 transformed is shown on the y-axis. Different constitutive promoters used to complement a *virE12* deletion mutant are shown as box and whisker plots with individual data points overlaid (n=64). Transformation by wild-type GV3101 and tobacco leaf without infiltration controls are shown. C) Proteomic spectral counts of VirE12 are shown when *virE12* is expressed from different constitutive synthetic promoters *in vitro* (n=3) Rows D-G show characterization of P_LacO_, P_TetR_, P_NahR_, P_JungleExpress_ respectively. From left to right Activity of inducible promoters driving RFP from pGingerBK plasmid backbone in tobacco (n=12), *Arabidopsis* (n=12), and *in vitro* (n=8). Inducer either not added (“None”), added at the half maximal induction concentration determined *in vitro* (“Mid”), or at the maximal induction concentration (“High”). The middle panel show the complementation of a *virE12* deletion by different inducible promoters as measured by transient GFP expression shown on the y-axis after log2 transformation(n=64). Inducer either not added (“None”), added at the half maximal induction concentration determined *in vitro* (“Mid”), or at the maximal induction concentration (“High”). The right panel shows proteomic spectral counts of VirE12 when expressed from different inducible promoters (n=3). Inducer either not added (“None”), added at the half maximal induction concentration determined *in vitro* (“Mid”), or at the maximal induction concentration (“High”).

Inducible promoters enable the dynamic control of gene expression strength, and thus could reduce the number of genetic designs needed to evaluate the impact of gene expression on AMT. Therefore, we then evaluated the expression of RFP from four inducible promoter systems (P_LacO_, P_TetR_, P_Jungle Express_, and P_NahR_) in culture media as well as in *N. benthamiana* and *A. thaliana* leaves, where the inducing compound was mixed with a bacterial suspension before infiltration into leaf tissue. While each of these systems displayed inducible expression in culture media (**Figure S2**), only P_LacO_ and P_TetR_ showed consistent inducibility in both host plant species (**Figures 1D**-**G****, Figure S3)**. Conversely, the P_Jungle Express_ promoter showed poor induction in both plant species, and P_NahR_ was expressed even in the absence of an added inducer within *A. thaliana* leaf tissue. These results demonstrate the importance of validating each promoter in its intended environment. For example, though functional *in vitro*, P_Jungle Express_ performed extremely poorly *in planta*. Similar results were observed for P_Jungle Express_, as it is possible the crystal violet inducer may be rapidly bound to plant tissue and thus not biologically available. Conversely, the salicylic acid inducer of P_NahR_ can be endogenously produced by plants as an immune response to pathogens such as *A. fabrum*, and thus may not be ideal for exerting orthogonal control of gene expression within different plants ^42^.

After testing all four promoter systems to complement a Δ*virE12* mutation, only the LacI inducible promoter, with the highest amount of added ligand tested, was able to recover transformation back to wild type levels (**Figure 1D**). As P_LacO_ showed the best plant orthogonality and ability to complement a *virE12* mutation, further designs requiring inducibility utilized the IPTG inducible promoter. While proteomics from cultures indicated that the levels of VirE2 expressed from inducible promoters were similar to the constitutive promoters (**Figures 1C-G**), the plant-specific utility of individual promoters suggests that they may be less useful for designing general functioning genetic circuits across plant environments.

### A quantitative understanding of the genetic contributions to AMT

To systematically assess the contributions individual *vir* genes have on plant transformation, we developed a quantitative virulence assay to measure the efficiency of T-DNA transfer into plant cells. To accomplish this, we first generated internal, in-frame deletion mutants of known functional non-regulatory *vir* gene clusters in *A. fabrum* GV3101: *virB1-11*, *virC12*, *virD12*, *virD3*, *virD4*, *virD5*, *virE12*, *virE3*, *virF*, *virH1*, *virH2*, and *virK* (**Figure 2A**). Using a transient GFP expression assay in *N. benthamiana* leaves, we observed that deletion of *virB1-11*, *virC12*, *virD12*, *virD4*, or *virE12* resulted in over 90% reduction in transformation efficiency (**Figures 2B**). Furthermore, loss of *virD5*, *virE3*, *virH1*, *virH2*, or *virK* significantly reduced transformation efficiency compared to wild-type, while deletion of *virD3* or *virF* showed no significant reduction in transformation efficiency (**Figure 2B**). Plasmid complementation of these deletions using the relatively weak promoter P_J23117_ restored wild-type transformation efficiencies in all deletion strains except *virB1-11*, *virD4*, *virD5*, and *virK* (**Figure 2B**). These results thus serve as a benchmark that, for the first time, allow for relative comparison of *vir* gene importance in AMT.

**Figure 2:**
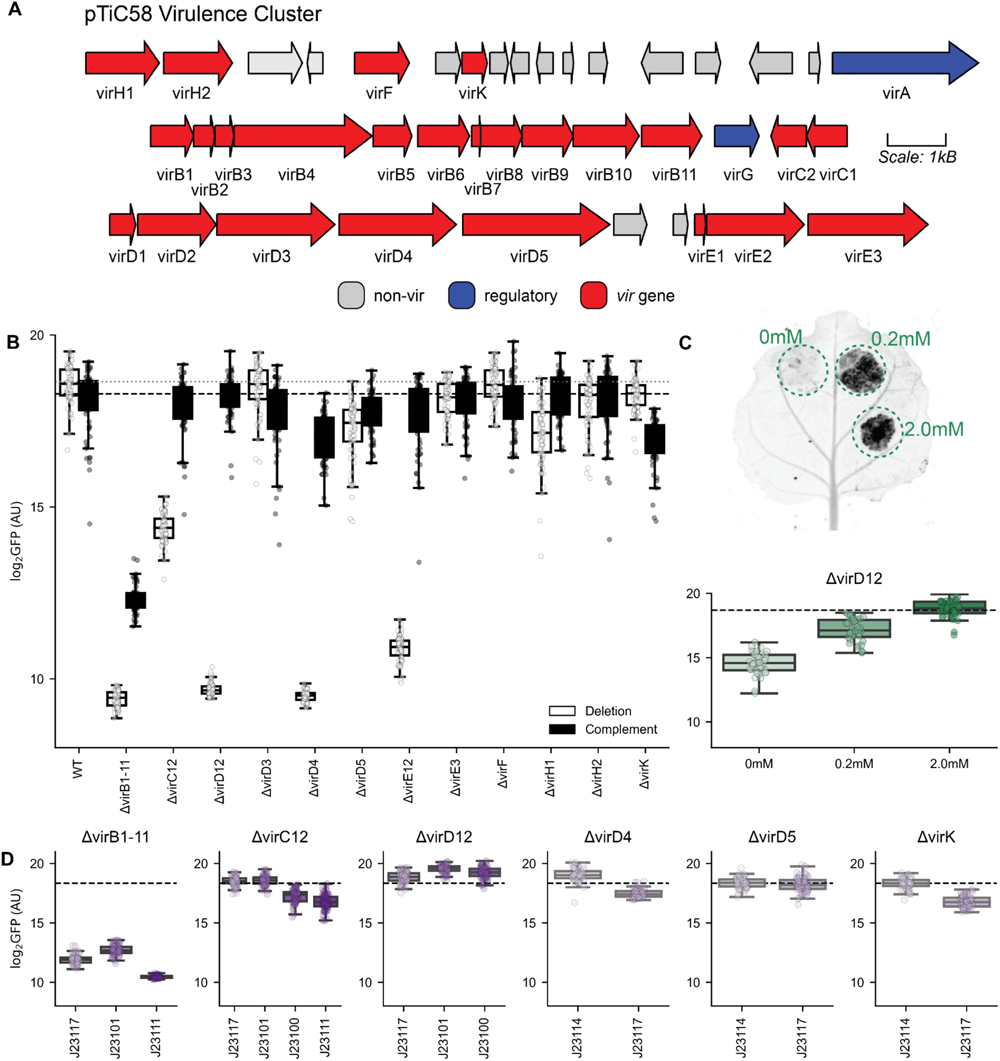
Quantitative assessment of *vir* gene impact on transformation: A) The *virulence* gene cluster from pTiC58. Known non-regulatory *vir* genes are shown in red, while regulatory *vir* genes are shown in blue. All other genes are grey. B) Effect of individual *vir* gene cluster deletion on tobacco transformation is measured by transient expression of GFP shown in log2 transformed AU in white, and complementation of the phenotype driven by P_J23117_ is shown in black (n=64). Dashed gray line shows transformation by wild-type GV3101, while dashed black line shows transformation by GV3101 expressing RFP from P_J23117_ as a control. C) Picture of a tobacco leaf expressing GFP delivered by a *virD12* deletion mutant complemented from an IPTG inducible promoter with varying levels of induction indicated. Below shows transient GFP expressed in tobacco when transformed by *virD12* deletion with different IPTG levels shown in log2 transformed AU is shown on the y-axis, with the concentration of IPTG used to induce the promoter on the x-axis (n=64). Dashed line shows wild-type level transformation D) Complementation of *vir* gene deletion mutants that showed trends from Figure S4 using different strength constitutive promoters. Transient tobacco-expressed GFP shown in log2 transformed AU is shown on the y-axis (n=64). Dashed line shows wild-type level transformation, colors of boxplots represent strength of constitutive promoters from Figure 1A.

To explore the effect of different expression levels on transformation efficiency, we then complemented each mutation with the inducible P_LacO_ promoter. Three phenotypes were observed: 1) the *virB1-11*, *virC12*, *virD12*, and *virE12* complementation strains had increasing transformation efficiency with increased induction; 2) the *virD3*, *virD4*, and *virD5* complementation strains had decreasing transformation efficiency as induction increased; and 3) the *virE3*, *virF*, *virH1*, *virH2*, and *virK* complementation strains showed no response to increasing induction (**Figure 2C, S4)**. Some of the relative decrease in transformation observed as *virD5* expression increases may be due to toxicity to the bacterium; however, similar toxicity was not observed with increased expression of *virD3* or *virD4* (**Figure S5)**. Previous work has shown that overexpression of *virD5* resulted in acute toxicity in eukaryotes, where it is localized to the nucleus, and may cause DNA damage^43,44^. Based on these results, we used constitutive promoters stronger or weaker than P_J23117_ to tune and optimize the expression of each *vir* gene cassette (**Figure 2D**). Strong expression of *virD12* improved transformation compared to wild-type by 135%. These results are in line with previous reports that overexpression of *virD12* improves transformation ^45^. Conversely, lower expression of *virD4* improved transformation 72% over wild-type. There was no significant improvement of transformation by increasing the expression of *virC12*, though expression from the stronger P_J23100_ and P_J23101_ promoters decreased transformation. Expression of *virD5* and *virK* from the weak P_J23114_ promoter was able to restore wild-type level transformation efficiency. Overall, these results demonstrate that transformation efficiency is highly sensitive to the expression strength of nearly all *vir* genes we evaluated, necessitating precise tuning for optimal DNA transfer.

Unlike other gene clusters, which were all complemented back to at least wild-type levels of transformation, we were only able to achieve ∼2% of wild-type transformation in a Δ*virB1-11* genetic background. Expressing *virB1-11* from the strong P_J23101_ promoter improved transformation over complementation using P_J23117_. However, complementation from the strongest promoter tested, P_J23111_, resulted in a significant reduction in transformation, suggesting that high-level expression may be toxic to the bacterium. The *virB* operon encodes the type 4 secretion system (T4SS), and previous studies genetically reconstructing secretion systems demonstrated the difficulty associated with engineering efficient transport ^11^., suggesting that engineering the T4SS may represent the bottleneck in engineering efforts.

In an attempt to improve *virB* complementation, we explored whether breaking the cluster into segments would improve our ability to complement the *virB* cluster. We knocked out *virB1-5* and *virB6-11* individually and attempted to complement these smaller mutations. Both of the smaller mutations predictably abolished transformation (**Figure S6A)**. Using the P_LacO_ inducible promoter, *virB1-5* showed a linear improvement of transformation with increased IPTG concentrations. However, *virB6-11* complementation plateaued at the median inducer concentration tested, with the highest level of induction causing a sharp decrease in transformation (**Figure S6B)**. The decrease in transformation is likely due to the extreme toxicity associated with *virB6-11* being expressed without the other T4SS genes, which greatly compromised growth (**Figure S6C)**. Constitutive complementation assays revealed the optimal promoters for complementing these deletions were the middle strength P_J23101_ for *virB1-5* which yielded ∼60% of wild-type transformation, and the relatively weak P_J23117_ for *virB6-11* which yielded ∼25% complementation (**Figure S6D)**. Based on this data we designed synthetic *virB1-11* complementation vectors that express *virB1-5* using three different promoters (weak-P_J23117_, medium-P_J23101_, and strong-P_J23100_), and *virB6-11* from the weak constitutive promoter, P_J23117_, with this cassette both downstream and upstream of *virB1-5* (**Figure S7A)**.

However, none of these vectors could complement as well as when *virB1-11* was expressed in its entirety. The vectors driving *virB1-5* from the strong P_J23100_ performed particularly poorly (**Figure S7A)**. To assess the performance of our synthetic complementation of *virB1-11* against the native P_virB_, we cloned the entire *virB1-11* operon in addition to its intergenic upstream and downstream DNA into a promoterless vector backbone. While this vector was able to complement transformation above the Δ*virB1-11* parent, it was still significantly less than both *virB1-11* expressed from P_LacO_, as well as *virB1-11* driven from P_J23101_ **(Figure S7B)**. These results may suggest that the *virB* cluster and other *vir* genes must be expressed from the same vector for efficient transformation.

### Screening natural diversity to assess the impact of disparate vir gene homologs on AMT

In many synthetically engineered metabolic pathways, multiple homologs of an enzyme are often evaluated to identify the optimal design needed to enhance flux towards the final product. To take a similar approach, we sampled the natural diversity of agrobacteria and then systematically tested homologs of non-regulatory *vir* genes for their ability to potentially improve transformation efficiency. It has recently been shown that at least 9 distinct lineages of pTi/pRi plasmids exist across the diversity of agrobacteria ^46,47^. (**Figure 3A**). To measure the effect allelic variation plays in plant transformation, we synthesized phylogenetically diverse alleles from each of the 9 pTi/pRi (**Table S1**) families and evaluated their ability to complement GV3101 deletion mutants of *virB1-11*, *virC12*, *virD12*, *virD4*, *virD5*, *virE12*, *virE3*, *virH1*, *virH2*, and *virF* in a tobacco transient expression system (**Figures 3B-K**).

**Figure 3:**
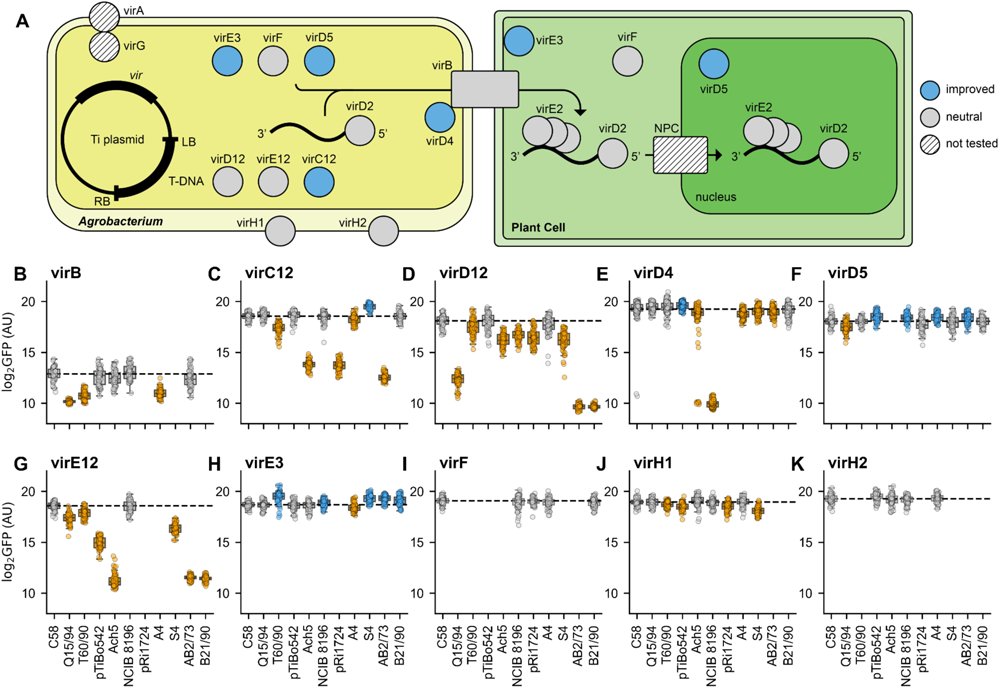
Impact of *vir* gene allele on tobacco transformation: A) Cartoon shows the localization of different *vir* gene products within the bacterial and plant cell during the AMT process. Gene products colored blue represent *vir* genes for homologs that outperformed wild-type in complementation assays. B-K) Plots show the effects of alleles of the indicated gene clusters in *A. fabrum* GV3101 deletion mutants complemented with constitutive promoters from a BBR1 origin plasmid. Box plots in yellow show alleles that are statistically worse than the wild-type allele, box plots in blue show alleles that are statistically superior than the wild-type allele, and box plots in white show alleles that are statistically indistinguishable from the wild-type allele. Statistical significance was determined using a Bonferroni corrected T-test (p-value < 0.05, n=64).

Of these clusters, we identified replacement alleles of *virC12* (91% improvement), *virD4* (13% improvement), *virD5* (35% improvement), and *virE3* (76% improvement) that resulted in improved complementation compared to the wild-type allele (C58). For *virD5*, 4 out of 9 alleles improved upon the wild type (**Figure 3F**). For *virE3*, 4 out of 9 tested also improved transformation compared to the native strain (**Figure 3H**). However, for the critical *vir* genes – *i.e.*, *virC12* (**Figure 3C**), *virD12* (**Figure 3D**), and *virE12* (**Figure 3G**) – the majority of homologs significantly reduced transformation. These results suggest that while homologs exist that can potentially improve transformation rates, AMT relies on multiple interactions between *vir* genes. Co-evolution between *vir* genes may limit the ability of distantly related homologs from functioning with one another.

To more specifically test whether phylogenetic distance from the wild-type allele impacts the ability for a *vir* gene to function in a non-native system, we correlated phylogenetic distance to the ability of a homolog to complement the C58 deletion mutant. With the exception of *virE12*, there were no significant correlations between phylogenetic distance and ability to complement (**Figure S8)**. Surprisingly, distantly related alleles were able to complement many of the *vir* gene mutants to the level of the wild-type allele. Many of the *vir* genes appear to be under purifying selection (d_N_/d_S_ < 1) across much of their coding sequence (**Figure S9)**. This selective pressure may keep critical residues needed for protein-protein interactions intact across evolutionary time, but further analysis will be required to identify whether such residues exist.

Given that we identified multiple homologs across 4 *vir* gene clusters that could improve transformation, we then asked if these homologs could be combined to further improve transformation. To this end, we generated a suite of plasmids, called pLoki, that contained either the critical genes *virC12*, *virD12*, *virD4*, and *virE12* (pLoki1) or these critical genes with the addition of *virD5* and *virE3* (pLoki2) **(Figure S10A-C)**. We constructed a total of 20 variants of both pLoki1 and pLoki2 that explored all possible combinations of both wild-type alleles (C58) and the alleles of *virC12*, *virD4*, *virD5*, and *virE3* that performed best from our initial screen under the control of promoters that optimally complemented deletions. These vectors were used to complement a deletion that spanned *virC2* to *virE3* in *A. fabrum* GV3101 (**Figure S10D)**.

The pLoki1 variant containing all C58 alleles restored ∼25% of wild-type transformation in a transient tobacco expression assay, whereas the pLoki2 variant containing all C58 alleles restored ∼65% (**Figure S10E)**. Across all pLoki variants, none that contained a non-native allele outperformed the pLoki plasmids that only contained wild-type genes (**Figure S10D**). Looking across pLoki variants, we observed that vectors containing *virD5* derived from pTiBo542 were significantly superior to those harboring the wild-type (**Figure S10F),** though the improvement was relatively minimal. Strains that contained *virD4* from pTiBo542 or *virE3* from pTiT60/94, however, were both worse than strains with the corresponding wild-type allele (**Figure S10G-H)**. Further understanding the molecular basis and evolutionary constraints in swapping *vir* genes may help direct future studies in harnessing the natural diversity of *vir* genes to improve AMT.

### Engineering a synthetic pTi enables orthogonal control of AMT

To exert predictable phenotypic control over AMT, the genotypic and regulatory makeup of a synthetic pTi must be composed of a defined set of genes controlled by promoters that are orthogonal to regulatory influence exerted by the plant environment. Based on our quantitative assessment of *vir* gene importance for tobacco transformation (**Figure 2**) and using optimal promoters previously identified (**Figure 1**), we first sought to identify the minimal set of genetic elements capable of plant transformation. Our initial design (pDimples0) contained a minimal set of essential *vir* genes (*i.e.*, *virB1-11*, *virD12,* and *virD4*) based on both our findings and previous work, which expressed the *virB* genes as a single operon controlled by P_LacO_, and the other genes controlled by optimally determined constitutive promoters (**Figure 4A**). This vector was then introduced into *A. fabrum* C58C1, a strain of *A. fabrum* which has been cured of its pTi, also harboring a binary vector expressing GFP on the T-DNA.

**Figure 4:**
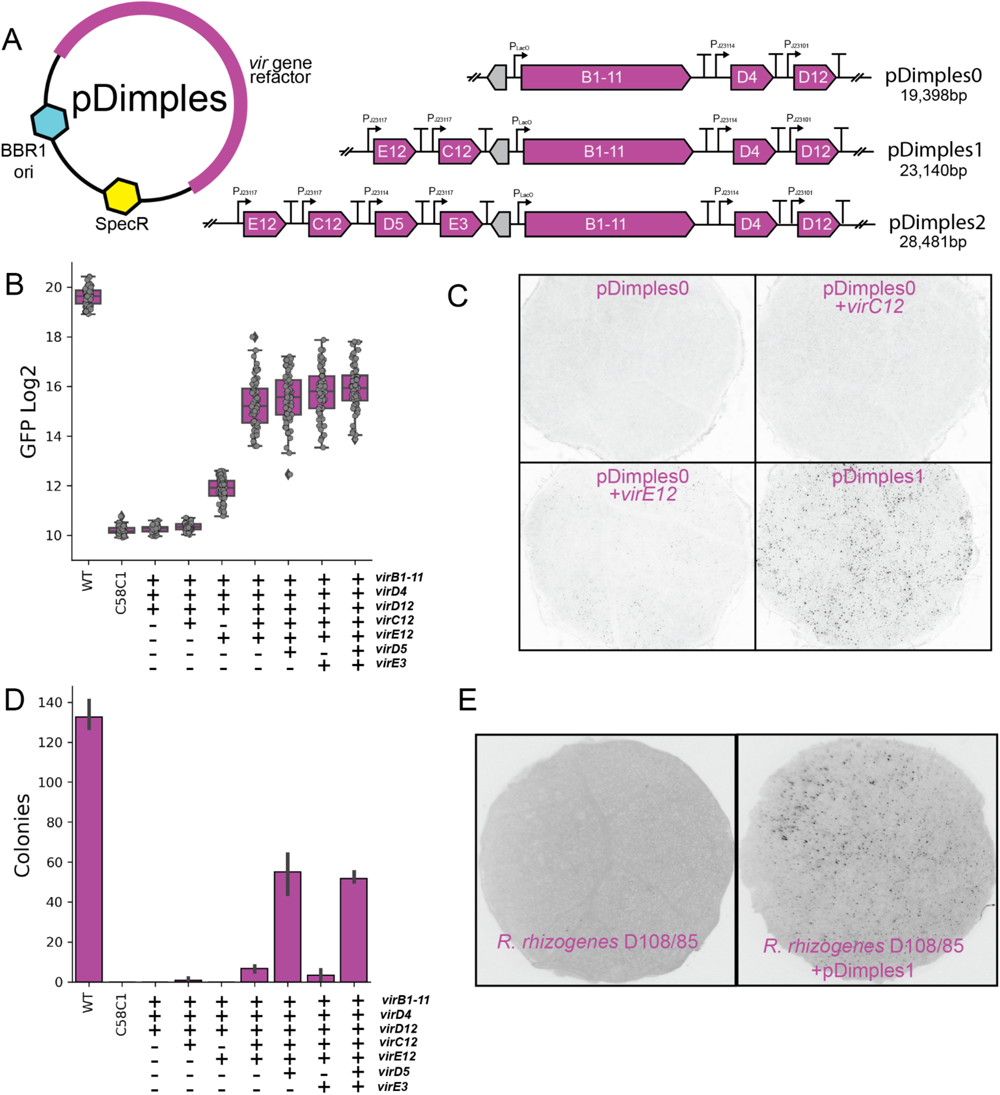
Synthetic refactoring of pTi: A) Genetic design of pDimples vectors. Variants of pDimples1 that only have either *virC12* or *virE12* (pDimples0.5), as well as variants of pDimples2 that have only virE3 or virD5 (pDimples1.5) were also constructed. B) GFP produced by transient transformation of tobacco leaves via synthetic pTi plasmids harbored in *A. fabrum* C58C1 (n=64). The Y-axis shows log2 transformed GFP (AU). C) Fluorescent microscopy of 6mM tobacco leaf punches infiltrated with *A. fabrum* C58C1 harboring minimal refactored pTi plasmids as well as a binary vector for the expression of a nuclear localized mScarlet. D) Transformation of *R. toruloides* by synthetically refactored pTi plasmids harbored in *A. fabrum* C58C. The average number of transformants obtained in three transformations is shown by refactored strains, as well as a wild-type strain of *A. fabrum* GV3101 and *A. fabrum* C58C1 harboring a binary vector but no refactored pTi. E) Fluorescent microscopy of 6mm tobacco leaf punches infiltrated with *R. rhizogenes* D108/85 harboring a binary vector for the expression of a nuclear localized mScarlet without (top) or with (bottom) pDimples1.0.

When this strain was introduced into tobacco leaves, there was no measurable increase in GFP signal when compared to leaves infiltrated with *A. fabrum* C58C1 carrying only the binary vector (**Figure 4B**). To further explore the minimal genetic requirements, we then generated two additional variants, which added either critical genes *virC12* (pDimples0.5-*virC12*) or *virE12* (pDimples0.5-*virE12*) upstream of the *virB* cluster. While pDimples0.5-*virC12* was unable to achieve any measurable tobacco transformation, pDimples0.5-*virE12* generated GFP above the control (**Figure 4B**). This finding was corroborated by experiments which expressed a nuclear localized mScarlet from the T-DNA. C58C1 containing the minimal pDimples0.5-*virE12* were able to form bright nuclear fluorescence in tobacco leaves, indicating successful T-DNA transfer into plant nuclei (**Figure 4C**). These findings experimentally define a minimal set of genes required for AMT of tobacco leaves and a starting point for rational engineering of synthetic pTi vectors.

To iterate upon and further optimize this design, we added both *virC12* and *virE12* upstream of the *virB* cluster (pDimples1.0) which dramatically improved transformation efficiency to 6.3% of wild type (**Figure 4B-C**). We then sought to evaluate whether the addition of either effector *vir* genes *virE3* or *virD5*, both of which significantly decreased transformation when deleted in *A. fabrum* GV3101, could improve transformation compared to pDimples1.0. Either gene was cloned in between *virC12* and *virB* clusters to generate pDimples1.5. While pDimples1.5-*virE3* improved transformation over pDimples1.0 to 8.3% of wild-type, pDimples1.5-*virD5* did not improve over pDimples1.0 (**Figure 4B**). When both *virE3* and *virD5* were added to create pDimples 2.0, transformation efficiency reached 9.1% of wild-type. While this was significantly improved from pDimples1.0, it was not significantly better than the addition of *virE3* alone (**Figure 4B**). These vectors were introduced into GV3101 with a deletion from *virA*-*virE3* constituting the majority of the *vir* genes and their essential positive regulators. When these complementation strains were compared to pDimples vectors harbored in C58C1, there was no significant difference in the transformation ability of strains with each vector (**Figure S11A)**. This suggests that – at least in the context of transient expression within tobacco – other genes on pTi may not play a significant role in the transformation process.

Attempts to optimize the expression of *virB* via complementation assays showed that P_LacO_ was an optimal choice to control the expression of the T4SS. The choice of the inducible P_LacO_ also allowed us to control the magnitude of transformation with the amount of IPTG that was co-infiltrated (**Figure S11B)**. As the ability of pDimples vectors to restore transformation was significantly less than that of the pLoki vectors (∼10% versus 75% restoration of wild-type *A. farbrum* GV3101 transformation), we concluded that suboptimal expression of the T4SS was likely a limiting factor. A possible bottleneck could be the availability of *virD4* which acts as a bridge between VirD2-conjugated T-DNA and the rest of the T4SS.

As *virD4* acts in concert with the T4SS to extrude the T-DNA, we sought to see if *virD4* expression limited transformation by replacing the very weak P_J23114_ promoter with the slightly stronger P_J23117_ promoter. However, this resulted in a significant decrease in transformation, indicating the bottleneck exists elsewhere (**Figure S11C)**. While no pDimples vector was able to restore wild-type level transformation to *A. fabrum* C58C1, pDimples1 and pDimples2 outperformed any attempt to complement a *virB1*-11 deletion. These results are consistent with our hypothesis that a specific ratio between the T4SS genes and other *vir* genes needs to be maintained for optimal transformation. Exploring this relationship further will likely be key in debottlenecking future engineering efforts.

Because *Agrobacterium* is also a critical tool for the transformation of many fungi ^48^, we evaluated the ability of the pDimples vectors to transform the oleaginous yeast *Rhodosporidium toruloides*. Unlike in tobacco, a small number of transformants were observed with pDimples0.5-*virC12* strains added, while no transformants were observed with pDimples0.5-*virE12* (**Figure 4D**). This is consistent with reports that *virE12* is not as important for fungal transformation as it is for plant transformation ^49^. While only 5% of wild type transformation efficiency was achieved with pDimples1.0, the addition of *virD5* dramatically increased transformation efficiency to 40% of wild-type (**Figure 4D**). This is intriguing because while VirD5 has been shown to localize to the nucleus of fungi ^43^, it was thought to be completely dispensable for fungal transformation ^49^. Contrary to previous thought, it is likely that *virD5* has a far more fundamental role in AMT than simply as a determinant of host range. Moreover, the addition of *virE3* by itself did not improve transformation, nor did it improve transformation efficiency when added in combination with *virD5*. As with tobacco experiments, fungal transformation was dependent on the presence of IPTG to induce *virB1-11* expression (**Figure S11D)**, with *R. toruloides* transformants being confirmed by colony PCR (**Figure S11E)**. Our synthetic pTi with differing minimal sets of *vir* genes demonstrate how a bottom-up engineering approach can define how AMT of fungi and plants differ, offering new opportunities to further dissect the contributory role of each *vir* gene in fungal AMT.

To test whether a synthetic pTi is sufficient to impart AMT outside of its native host context, we sought to test our engineered designs in a bacterium beyond *A. fabrum*. To this end we introduced pDimples1.0 into *Rhizobium rhizogenes* D108/85, a non-pathogenic strain without a native pRi or pTi plasmid and diverged from *A. fabrum* ∼200 million years ago ^46^. When *R. rhizogenes* was infiltrated into tobacco leaves carrying a binary vector expressing nuclear-localized mScarlet, no red nuclei were observed (**Figure 4E**). Yet with the addition of pDimples1.0, red nuclei were observed that produced significantly more fluorescent signal than the parent strain (**Figure 4E, Figure S11F)**. Together, our results demonstrate that the design and construction of synthetic pTi can be used to: 1) identify the core set of genes that are necessary and sufficient for AMT, 2) describe the contributory role of accessory *vir* genes, 3) divorce AMT from its native regulation, and 4) transfer this complex trait into other bacteria.

## Conclusion

Here, we leveraged a comprehensive and quantitative understanding of each *vir* gene cluster to build synthetic pTi plasmids that define the minimal transferable set required for AMT of both plants and fungi. Optimization of this set will allow us to better understand host-specificity between natural strains of *Agrobacterium* and to engineer laboratory strains with superior transformation properties. Furthermore, our analysis of how allelic variation of *vir* genes impacts transformation suggests there are likely untapped genetic resources to improve AMT. Overall, this work will also serve to guide related research studying host-microbe interactions, specifically those of plant-associated bacteria. For example, recent research that developed minimized versions of the nitrogen fixing pSymA in the root nodule-associated legume symbiont *Sinorhizobium meliloti* could be furthered by evaluating the impact of gene expression on individual genes ^13^.

We compared bacterial synthetic biology parts both *in vitro* and *in planta*, revealing that while constitutive synthetic promoters will likely perform similarly in different environments, the performance of inducible systems may be highly variable. Further characterization of synthetic regulatory elements *in situ* will enable more precise engineering. However, by using these tools to replace the master regulatory VirA/G system with synthetic regulation, we not only gained precise control of individual gene expression, but also insulated the bacteria from host mechanisms that interfere with gene expression, which has been previously observed ^50,51^. In fungi, current methods require long *vir* gene induction times in conditions that may not be optimal for the growth of certain fungi, which could be bypassed using synthetic pTi ^49,52^. Thus, separating AMT induction from its native inducing conditions (*i.e.*, low pH, sugar, and phenolic compounds) may also provide unique opportunities in improving fungal transformations.

By mobilizing the transformation phenotype via pDimples into *R. rhizogenes*, we open the door to another promising avenue of AMT engineering: transferring the complex *vir* machinery to other bacteria. As *A. fabrum* can elicit plant immunity that impede transformation, multiple efforts have been made recently to circumvent this either through mutation of known immunogenic loci ^53^ or the addition of immune suppressing systems ^54^. This work lays the foundation to developing synthetic pTi that function in bacteria that elicit minimal immune responses across plant species, potentially enabling the transformation of those that have traditionally been recalcitrant to genetic modification.

In synthetic biology, our inability to efficiently transform new organisms represents the biggest bottleneck to dramatically expanding the scope and range of species that can be utilized. Given the wide diversity of eukaryotes that can be transformed by *Agrobacterium*, future synthetic pTi may be optimized to target currently untransformable organisms and enable entirely new areas of biotechnology.

## Materials and Methods

### Media, chemicals, and culture conditions

Routine bacterial cultures were grown in Luria-Bertani (LB) Miller medium (BD Biosciences, USA). *E. coli* was grown at 37 °C, while *A. fabrum* was grown at 30 °C unless otherwise noted. Cultures were supplemented with kanamycin (50 mg/L, Sigma Aldrich, USA), gentamicin (30 mg/L, Fisher Scientific, USA), or spectinomycin (100mg/L, Sigma Aldrich, USA), when indicated. All other compounds unless otherwise specified were purchased through Sigma Aldrich. Bacterial kinetic growth curves were performed as described previously ^40^.

### Strains and plasmids

All bacterial strains and plasmids used in this work are listed in **Supplemental Table 1 and 2**. All strains and plasmids created in this work are viewable through the public instance of the JBEI registry. (https://public-registry.jbei.org/folders/814). All plasmids generated in this paper were designed using Device Editor and Vector Editor software, while all primers used for the construction of plasmids were designed using j5 software ^55–57^. Synthetic DNA was synthesized from Twist Biosciences. Plasmids were assembled via Gibson Assembly using standard protocols ^58^, Golden Gate Assembly using standard protocols ^59^, or restriction digest followed by ligation with T4 ligase as previously described ^60^. Plasmids were routinely isolated using the Qiaprep Spin Miniprep kit (Qiagen, USA), and all primers were purchased from Integrated DNA Technologies (IDT, Coralville, IA). Plasmid sequences were verified using whole plasmid sequencing (Primordium Labs, Monrovia, CA). *Agrobacterium* was routinely transformed via electroporation as described previously ^61^.

### Construction of deletion mutants

Deletion mutants in *A. fabrum* GV3101 were constructed by homologous recombination and *sacB* counterselection using the allelic exchange as described previously ^62^. Briefly, homology fragments of 1 kbp up-and downstream of the target gene, including the start and stop codons respectively, were cloned into pMQ30K - a kanamycin resistance-bearing derivative of pMQ30 ^63^. Plasmids were then transformed via electroporation into *E. coli* S17 and then mated into *A. fabrum* via conjugation. Transconjugants were selected for on LB Agar plates supplemented with kanamycin 50 mg/mL, and rifampicin 100 mg/mL. Transconjugants were then grown overnight on LB media also supplemented with 50 mg/mL kanamycin, and 100 mg/mL rifampicin, and then plated on LB Agar with no NaCl supplemented with 10% w/v sucrose. Putative deletions were restreaked on LB Agar with no NaCl supplemented with 10% w/v sucrose, and then were screened via PCR with primers flanking the target gene to confirm gene deletion.

### Synthetic part characterization

Characterization of pGinger vectors harbored by *A. fabrum in vitro* was performed as previously described for other bacteria ^40^. Briefly, *A. fabrum* C58C1 with different pGinger vectors were grown overnight in 10mL of LB supplemented with kanamycin overnight at 30°C with 250 rpm shaking and then diluted 1:100 into 500 μL of fresh LB media with kanamycin in a deep-well 96-well plate (Corning) For inducible promoters, chemical inducers were added in two-fold dilutions before incubation. Cells were then grown at 30°C for 24-hours while shaking at 250 rpm, and then 100 μL was measured for absorbance at OD_600_ as well as for RFP fluorescence using an excitation wavelength of 590 nm and an emission wavelength of 635 nm with a gain setting of 75 on a BioTek Synergy H1 microplate reader (Agilent).

To evaluate the performance of synthetic promoters *in planta,* strains were grown in 5mL LB media with kanamycin at 30°C with 250 rpm shaking overnight, and then diluted 1:5 with fresh media then grown for an additional 3 hours at 30°C with 250 rpm shaking. Cultures were then adjusted to an absorbance at OD_600_ of 1.0 in agroinfiltration buffer (10mM MgCl_2_, 10mM MES, 200µM acetosyringone, pH 5.6), and infiltrated into either *N. benthamiana* or *A. thaliana* leaf tissue. When appropriate chemical inducers were added to the agroinfiltration media immediately before leaf infiltration. Either one, or three days post-infiltration 6mm leaf disks were excised from each Agro-infiltrated leaf using a hole puncher and placed atop 300µL of water in a black, clear-bottom, 96-well microtiter plate (Corning). GFP fluorescence of each leaf disk was then measured using a BioTek Synergy H1 microplate reader (Agilent) with an excitation wavelength of 488 nm and measurement wavelength of 520 nm.

### Plant Growth Conditions

*A. thaliana* were germinated and grown in Sunshine Mix #1 soil (Sungro) in a Percival growth chamber at 22°C and 60% humidity using a 8/16 hour light/dark cycle with a daytime PPFD of ∼200 µmol/m^2^s. *N. benthamiana* plants were grown according to a previously described standardized lab protocol ^41^. All tobacco growth was conducted in an indoor growth room at 25°C and 60% humidity using a 16/8 hour light/dark cycle with a daytime PPFD of ∼120 µmol/m^2^s. Plants were maintained in Sunshine Mix #4 soil (Sungro) supplemented with Osmocote 14-14-14 fertilizer (ICL) at 5mL/L and agroinfiltrated 29 days after seed sowing.

### Tobacco Infiltration and Leaf Punch Assay

*A. fabrum* strains were grown in LB liquid media containing necessary antibiotics (50 µg/mL rifampicin, 30 µg/mL gentamicin, 50 µg/mL kanamycin, and 100 µg/mL spectinomycin for most strains) to an OD600 between 0.6 and 1.0 before pelleting. Cells were then prepared for infiltration by resuspension in agroinfiltration buffer (10mM MgCl_2_, 10mM MES, 200µM acetosyringone, pH 5.6) to a final OD600 of 1.0 and were allowed to induce for 2 hours in infiltration buffer at room temperature. When appropriate, chemical inducers (i.e. IPTG) were added during the 2 hour induction period. Each strain was then infiltrated into the fourth and fifth leaf (counting down from the top) of eight biological replicate tobacco plants. GFP transgene expression in agroinfiltrated leaves was then assessed by a leaf disk fluorescence assay three days post-infiltration. Four 6mm leaf disks were excised from each agroinfiltrated leaf using a hole puncher and placed atop 300µL of water in a black, clear-bottom, 96-well microtiter plate (Corning). GFP fluorescence of each leaf disk was then measured using a BioTek Synergy H1 microplate reader (Agilent) with an excitation wavelength of 488 nm and measurement wavelength of 520 nm.

### Rhodospordium toruloides Transformation

*Agrobacterium tumefaciens* mediated transformation was performed on *Rhodosporidium toruloides* IFO0880 with a codon optimized epi-isozizaene synthase from *Streptomyces coelicolor* A3(2) (JPUB_013523) ^64^ as previously described ^65^. When appropriate 2mM IPTG was added to agrobacterium induction media. Transformants were confirmed via colony PCR specific to the integrated T-DNA.

### Proteomic Analysis

Proteins from *A. fabrum* samples were extracted using a previously described chloroform/methanol precipitation method ^66^. Extracted proteins were resuspended in the 100 mM ammonium bicarbonate buffer supplemented with 20% methanol, and protein concentration was determined by the DC assay (BioRad). Protein reduction was accomplished using 5 mM tris 2-(carboxyethyl)phosphine (TCEP) for 30 min at room temperature, and alkylation was performed with 10 mM iodoacetamide (IAM; final concentration) for 30 min at room temperature in the dark. Overnight digestion with trypsin was accomplished with a 1:50 trypsin:total protein ratio. The resulting peptide samples were analyzed on an Agilent 1290 UHPLC system coupled to a Thermo scientific Obitrap Exploris 480 mass spectrometer for discovery proteomics ^67^. Briefly, 20 µg of tryptic peptides were loaded onto an Ascentis® (Sigma–Aldrich) ES-C18 column (2.1 mm × 100 mm, 2.7 μm particle size, operated at 60°C) and were eluted from the column by using a 10 minute gradient from 98% buffer A (0.1 % FA in H2O) and 2% buffer B (0.1% FA in acetonitrile) to 65% buffer A and 35% buffer B. The eluting peptides were introduced to the mass spectrometer operating in positive-ion mode. Full MS survey scans were acquired in the range of 300-1200 m/z at 60,000 resolution. The automatic gain control (AGC) target was set at 3e6 and the maximum injection time was set to 60 ms. Top 10 multiply charged precursor ions (2-5) were isolated for higher-energy collisional dissociation (HCD) MS/MS using a 1.6 m/z isolation window and were accumulated until they either reached an AGC target value of 1e5 or a maximum injection time of 50 ms. MS/MS data were generated with a normalized collision energy (NCE) of 30, at a resolution of 15,000. Upon fragmentation precursor ions were dynamically excluded for 10 s after the first fragmentation event. The acquired LCMS raw data were converted to mgf files and searched against the latest uniprot *A. tumefaciens* protein database with Mascot search engine version 2.3.02 (Matrix Science). The resulting search results were filtered and analyzed by Scaffold v 5.0 (Proteome Software Inc.). The normalized spectra count of identified proteins were exported for relative quantitative analysis.

### Bioinformatic Analyses

Sequences of individual *vir* genes from genomes of all sequenced *Agrobacterium* were identified and extracted as previously described ^68^. MACSE v. 2.07 with the parameter “-prog alignSequences” was used to generate codon alignments for each *vir* gene dataset ^69^. The HYPHY v2.2 program “cln” was used to remove identical sequences and stop codons from each alignment ^70^. IQ-TREE v. 1.6.12 with the default parameters was used to generate a phylogeny for each dataset ^71^. The HYPHY program FUBAR with the codon alignment, phylogeny, and a probability threshold of 0.9 was used to calculate per-site d_N_/d_S_ and detect signals of positive or purifying selection.

### Statistical analyses and data presentation

All numerical data were analyzed using custom Python scripts. All graphs were visualized using either Seaborn or Matplotlib ^72,73^. Calculation of 95% confidence intervals, standard deviations, and T-test statistics were conducted via the Scipy library ^74^. Bonferroni corrections were calculated using the MNE python library ^75^.

Alleles of homologous *vir* genes were aligned using MAFFT v. 7.508 ^76^ and converted into phylogenetic trees using FastTree v. 2.1.11 ^77^. Phylogenetic distance was calculated using dendropy v. 4.6.1 ^78^.

## Acknowledgements

We would like to thank Catharine Adams, Adam Arkin, and William Moore for helpful discussions during the preparation of this manuscript. We would like to thank Sasilada Sirirungruang and Simon Alamos for help with plant growth and microscopy. We would also like to thank Rachel Li, Nick Harris, Charles Denby, and Jutta Dalton for their support during the Covid-19 pandemic. *A. fabrum* C58C1 was received from John Zupan at UC Berkeley. MGT is a Simons Foundation Awardee of the Life Sciences Research Foundation. LMW is funded through the National Science Foundation Graduate Research Fellowship. AJW was funded in part by startup funding from the Department of Botany and Plant Pathology at Oregon State University. JHC is supported in part by the National Institute of Food and Agriculture, US Department of Agriculture (2022-67013-36883). Synthesis of *vir* gene alleles was supported by the JGI BRC proposal WIP# 507140: Synthetic minimal redesign of plant transformation plasmids. This work was part of the DOE Joint BioEnergy Institute (https://www.jbei.org) supported by the U. S. Department of Energy, Office of Science, Office of Biological and Environmental Research, supported by the U.S. Department of Energy, Energy Efficiency and Renewable Energy, Bioenergy Technologies Office, through contract DE-AC02-05CH11231 between Lawrence Berkeley National Laboratory and the U.S. Department of Energy. The funders had no role in manuscript preparation or the decision to publish. The views and opinions of the authors expressed herein do not necessarily state or reflect those of the United States Government or any agency thereof. Neither the United States Government nor any agency thereof, nor any of their employees, makes any warranty, expressed or implied, or assumes any legal liability or responsibility for the accuracy, completeness, or usefulness of any information, apparatus, product, or process disclosed, or represents that its use would not infringe privately owned rights. The United States Government retains and the publisher, by accepting the article for publication, acknowledges that the United States Government retains a nonexclusive, paid-up, irrevocable, worldwide license to publish or reproduce the published form of this manuscript, or allow others to do so, for United States Government purposes. The Department of Energy will provide public access to these results of federally sponsored research in accordance with the DOE Public Access Plan (http://energy.gov/downloads/doe-public-access-plan).

## Contributions

Conceptualization, M.G.T., P.M.S.; Methodology, M.G.T.,L.D.K, A.N.P., L.W., A.W., G.M.G.; Investigation, M.G.T., L.D.K., G.M.G., L.M. W., A.N.P., M.S., K.V., S.S., K.M., S.A, N.F.C.H, D.S., C.T., R.C., S.L., J.C., H.P., J.W.G, Y.C., A.J.W.; Writing – Original Draft, M.G.T.; Writing – Review and Editing, All authors.; Resources and supervision, D.L., C.J.P., J.M.G., H.V.S A.W., J.H.C., J.D.K., P.M.S.

