## Supplementary material for "Genetically refactored *Agrobacterium*-mediated transformation": Supplmental Figures

**Supplementary Figures**

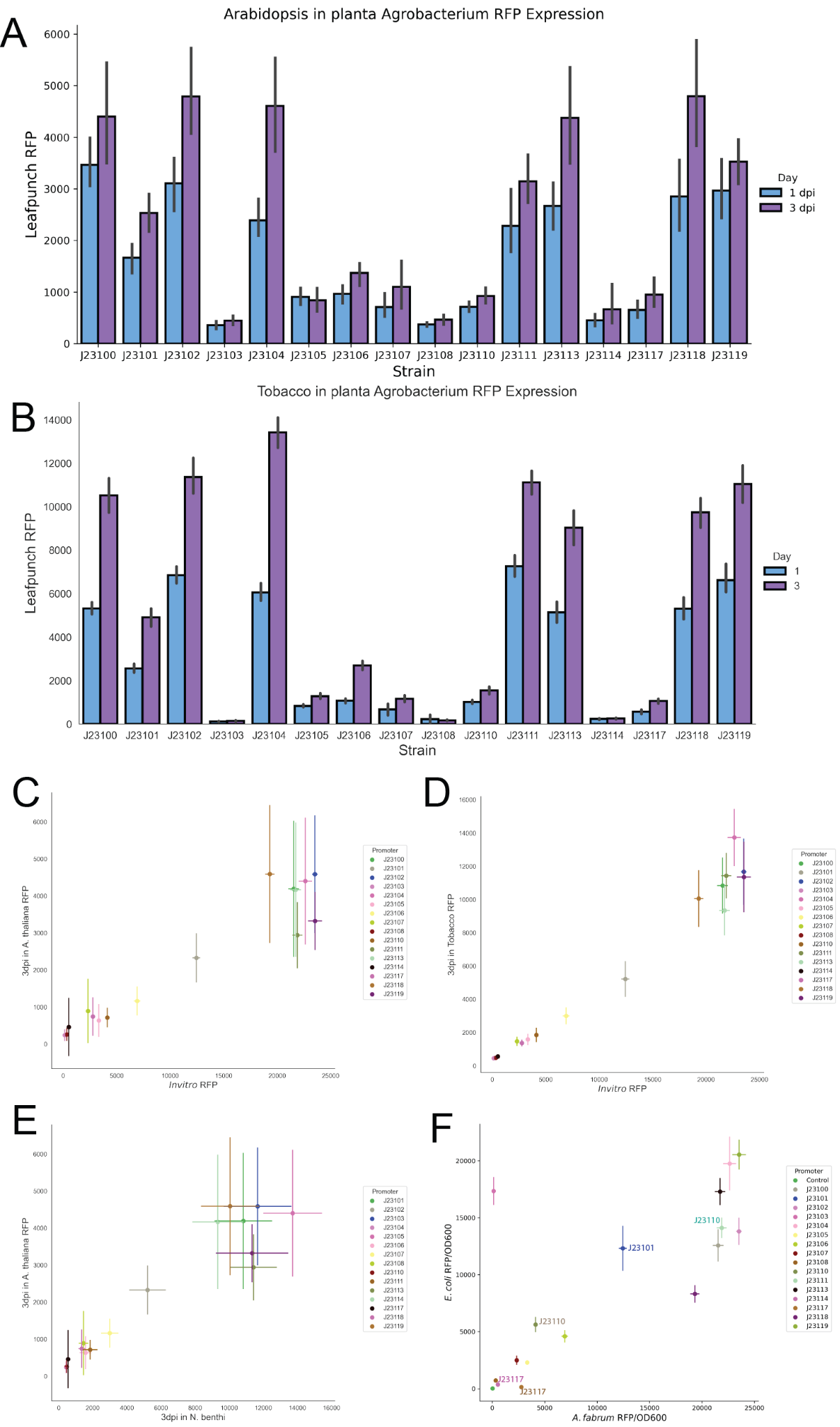

**Figure S1**: Characterization of constitutive promoters: A) RFP expression from pGingerBK constitutive promoters within *A. fabrum* after infiltration into *A. thaliana* leaves 1 or 3 days post. Error bars show standard deviations (n=12). B) RFP expression from pGingerBK constitutive promoters within *A. fabrum* after infiltration into tobacco leaves 1 or 3 days post. Error bars show standard deviations (n=12). C) Scatterplot shows correlation between *in vitro* expression of constitutive promoters and expression within *A. thaliana* leaf tissue 3 days post infiltration. D) Scatterplot shows correlation between *in vitro* expression of constitutive promoters and expression within tobacco leaf tissue 3 days post infiltration E) Scatterplot shows correlation between expression of constitutive promoters within tobacco leaf tissue 3 days post infiltration and expression within *A. thaliana* leaf tissue 3 days post infiltration. F) Scatterplot shows correlation of RFP expression from pGingerBK constitutive promoters in *E. coli* and *A. fabrum in vitro*. Promoters used in this study are highlighted. Error bars show standard deviations (n=8).

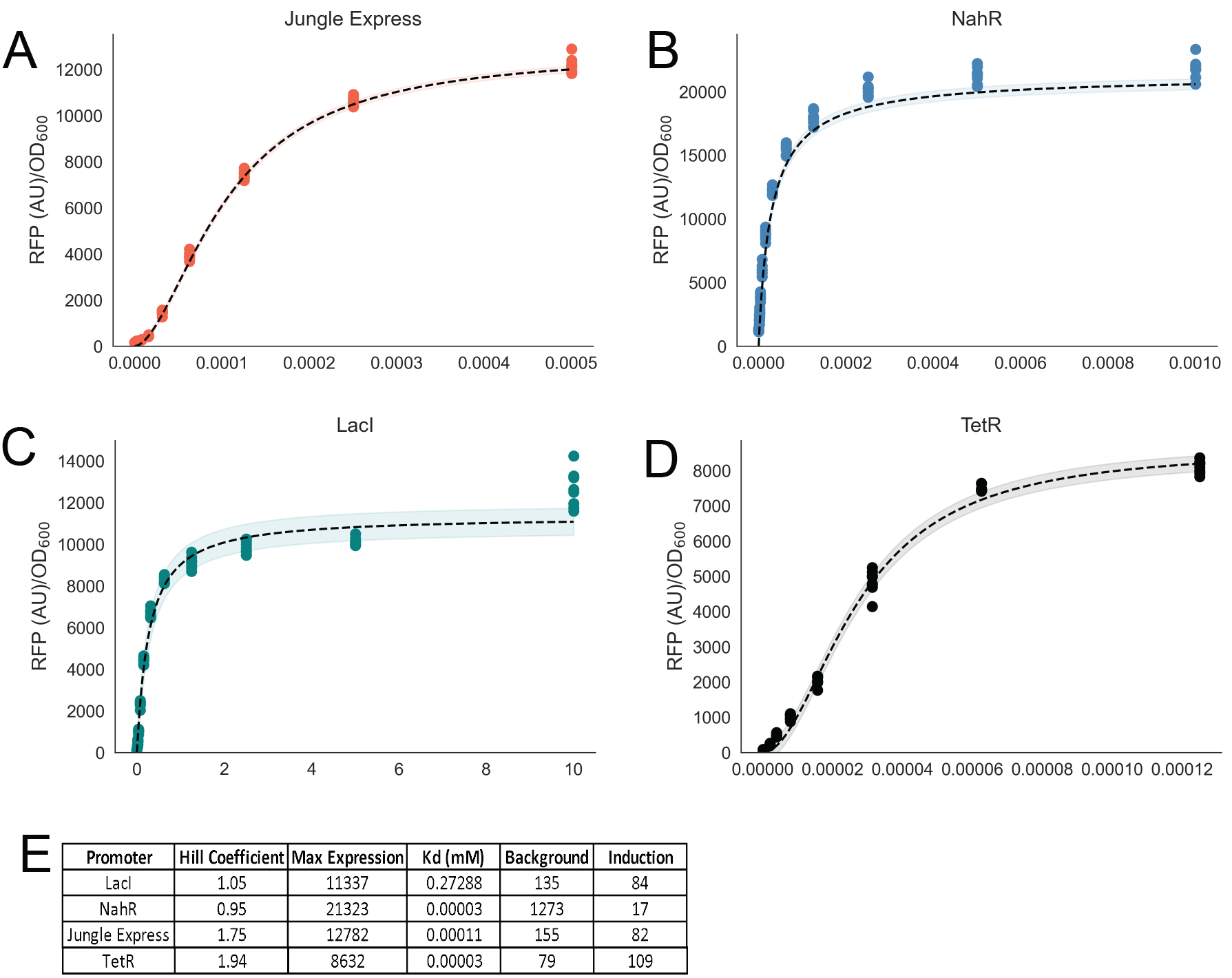

**Figure S2:** B) Dose response curves of the four inducible promoters used in this study. Graphs show RFP expression of A) Jungle Express B) NahR C) LacO and D) TetR (n=8). Dashed lines show fit to the Hill equation, and shaded area represents the confidence of the fit. E) Table shows relevant parameters of the Hill Equation fit.

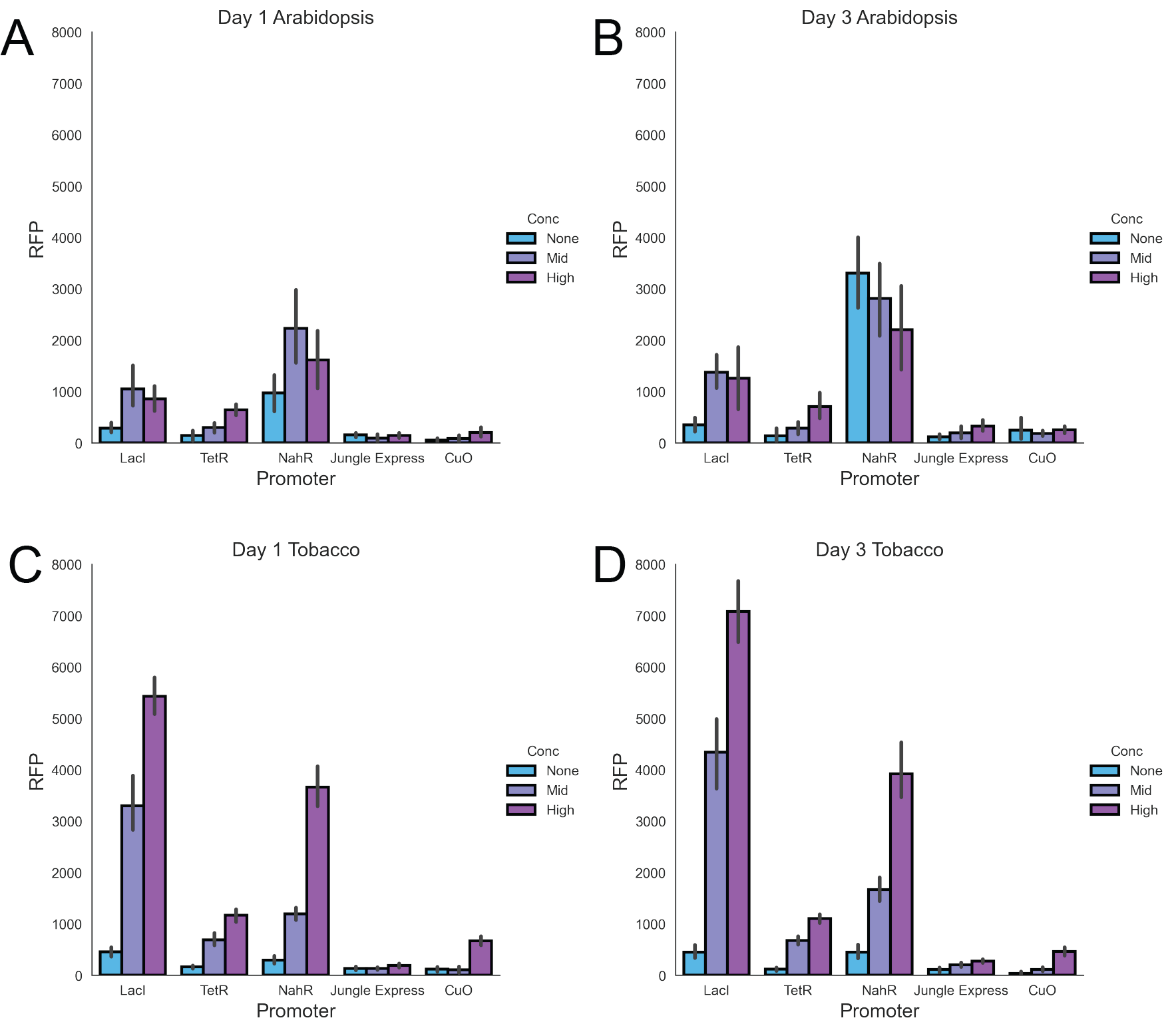

**Figure S3**: Graphs show expression of *A. fabrum* C58C1 RFP from inducible promoters within *A. thaliana* 1 (A) or 3 (B) days post infiltration, and tobacco 1 (C) or 3 (D) days post infiltration, at different levels of promoter induction (n=12).

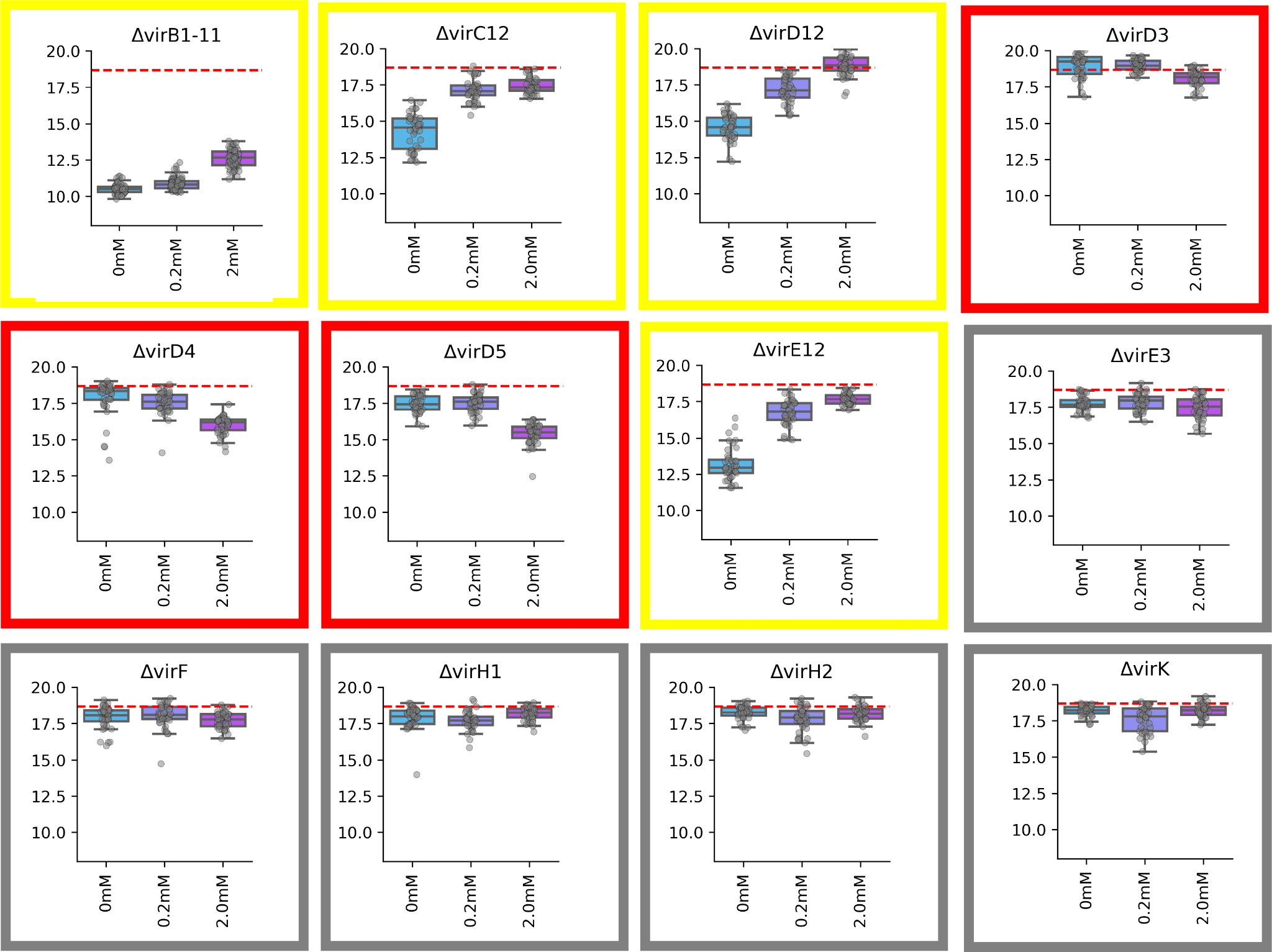

**Figure S4:** Impact of increasing *vir* gene expression on tobacco transient transformation. Graphs show complementation of *vir* gene cluster deletions of *A. fabrum* GV3101 with the IPTG inducible pGingerBS-LacO plasmid. The y-axis shows transient GFP production and the x-axis shows complementation at different levels of IPTG induction (n=64). Plots outlined in yellow show a positive correlation between induction and transformation, plots outlined in red show a negative correlation between induction and transformation, and plots outlined in gray show no correlation between induction and transformation. Dashed red lines on plots show a wild-type level of transformation.

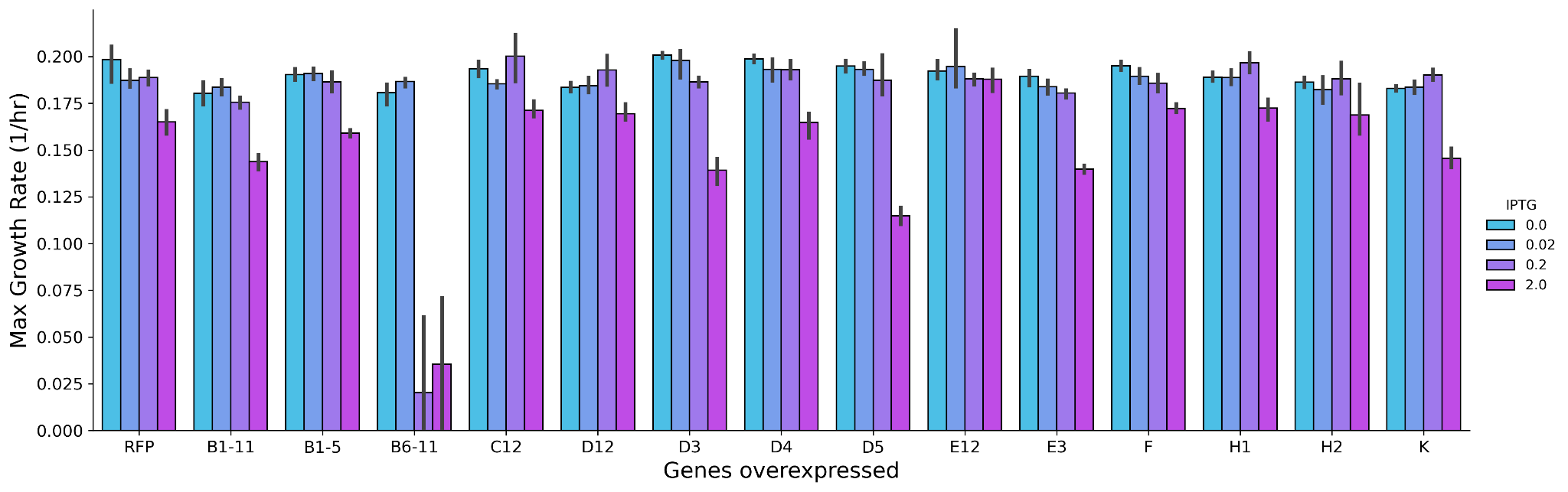

**Figure S5:** Growth rate of *A. fabrum* GV3101 *vir* gene cluster deletion strains complemented with IPTG inducible pGingerBS-LacO plasmid. Graph shows the maximal logarithmic phase growth rate of strains grown in a microplate reader at different levels of IPTG induction (n=4). GV3101 wild-type expressing RFP is shown as a control. Error bars represent standard deviations.

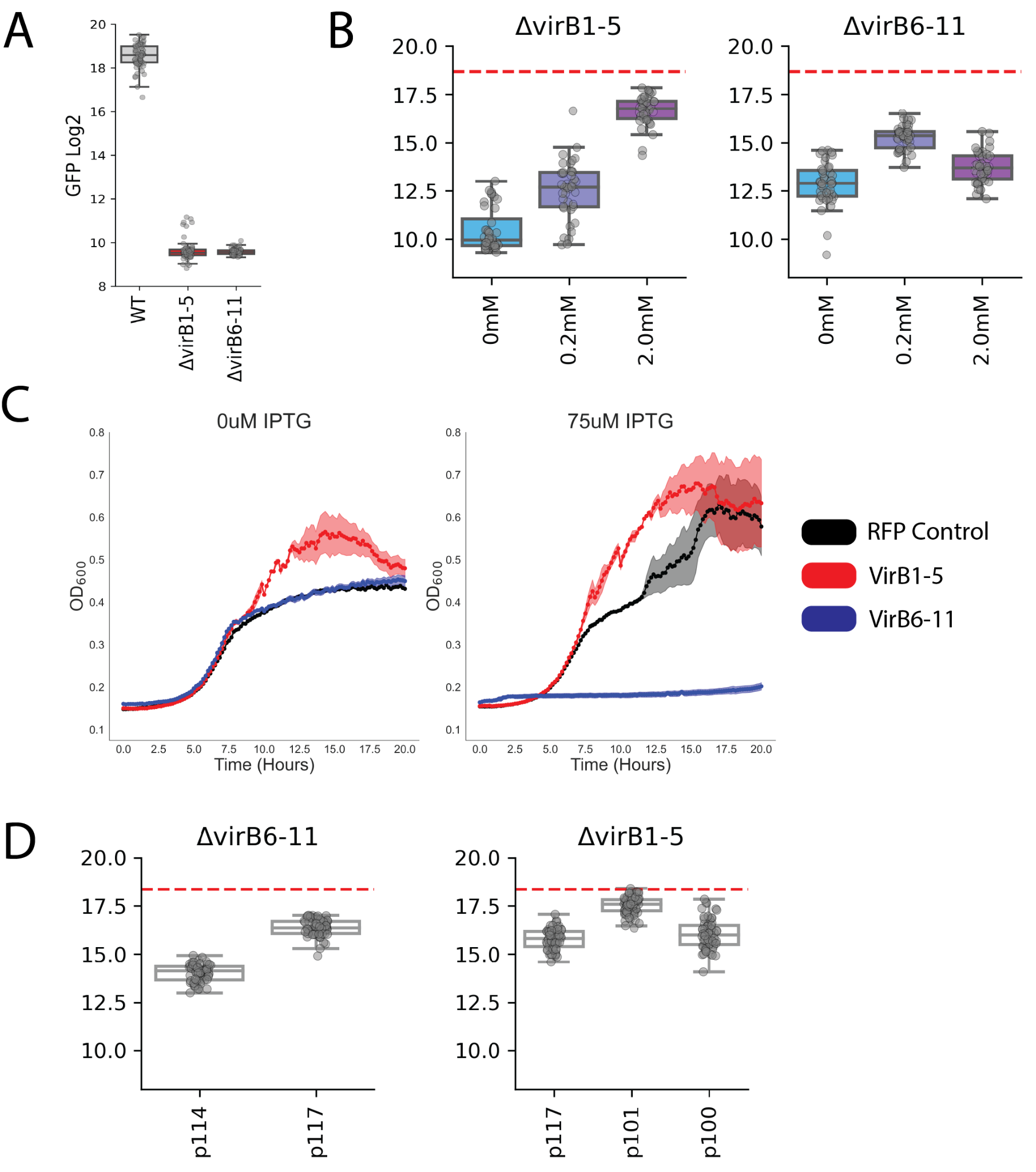

**Figure S6:** Complementing *virB1-5* and *virB6-11* deletion mutants: A) Effect of knocking out either *virB1-5* or *virB6-11* is measured by transient expression of GFP shown in log2 transformed AU (n=64). B) Graphs show complementation of either *virB1-5* or *virB6-11* gene cluster deletions of *A. fabrum* GV3101 with the IPTG inducible pGingerBS-LacO plasmid. The y-axis shows transient GFP production and the x-axis shows complementation at different levels of IPTG induction (n=64). Dashed red lines on plots show a wild-type level of transformation. C) Growth of wild-type *A. fabrum* GV3101 expressing RFP from IPTG inducible pGingerBS-LacO plasmid (black), a *virB1-5* deletion mutant expressing *virB1-5* from an IPTG inducible pGingerBS-LacO plasmid (red), and a *virB6-11* deletion mutant expressing *virB6-11* from an IPTG inducible pGingerBS-LacO plasmid (red), with either 0 (left panel) or 75uM (right panel) IPTG added (n=4). D) Evaluating constitutive promoter complementation of either *virB1-5* or *virB6-11* deletion mutants of *A. fabrum* GV3101. The y-axis shows log2 transformed GFP (AU) from transient tobacco infiltrations, x-axis represents different constitutive promoters tested (n=64). Dashed red lines on plots show a wild-type level of transformation.

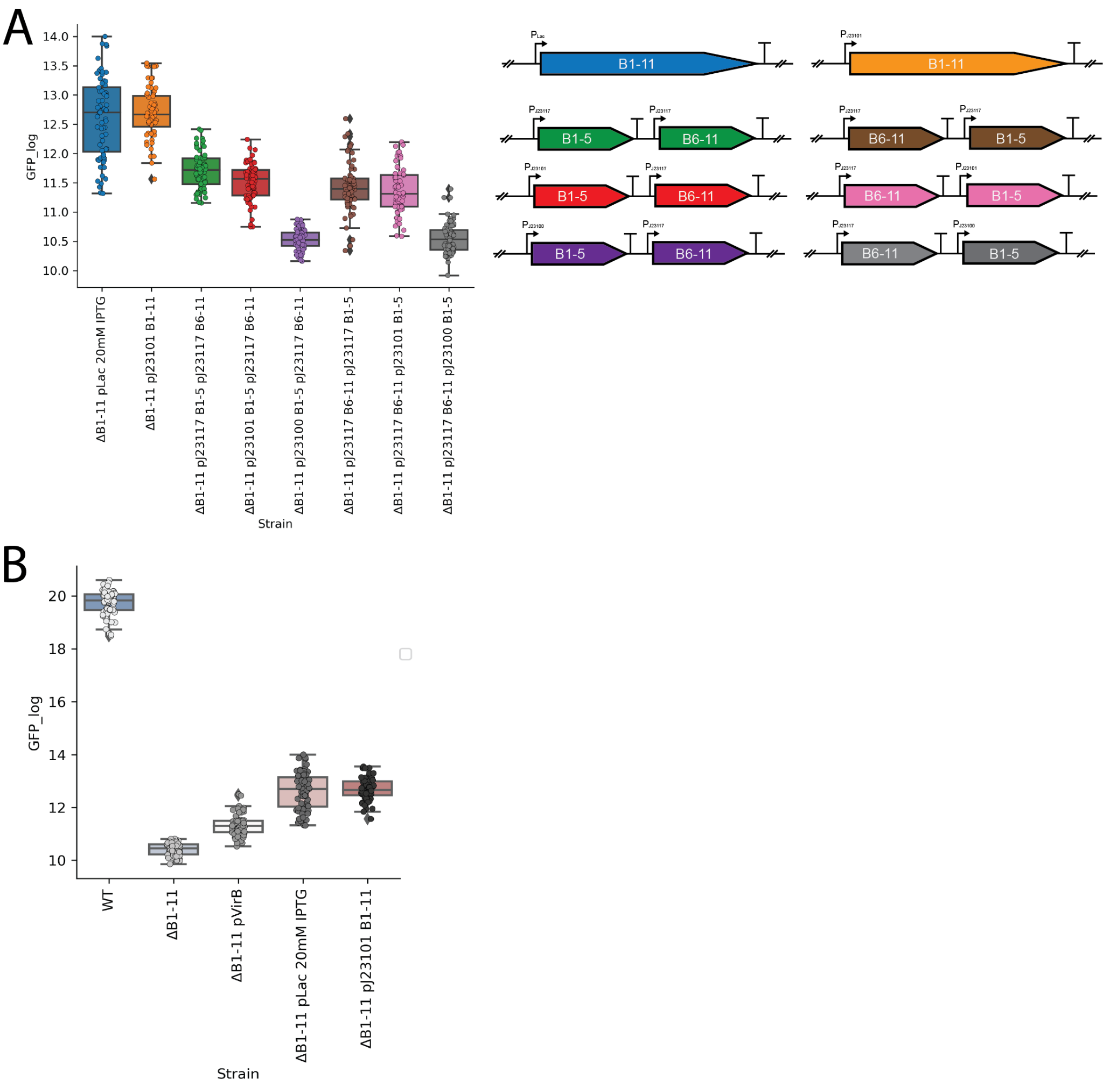

**Figure S7:** Optimization of *virB1-11* complementation: A) Evaluating different complementation vectors of a *virB1-11* deletion mutant of *A. fabrum* GV3101. The y-axis shows log2 transformed GFP (AU) from transient tobacco infiltrations, x-axis represents different constitutive promoters tested (n=64). Genetic diagrams to the right are color coordinated with the boxplots to the left. B) Comparing P_virB_, P_LacO_, and P_J23101_ complementation vectors of a *virB1-11* deletion mutant of *A. fabrum* GV3101. The y-axis shows log2 transformed GFP (AU) from transient tobacco infiltrations, x-axis represents promoters tested (n=64).

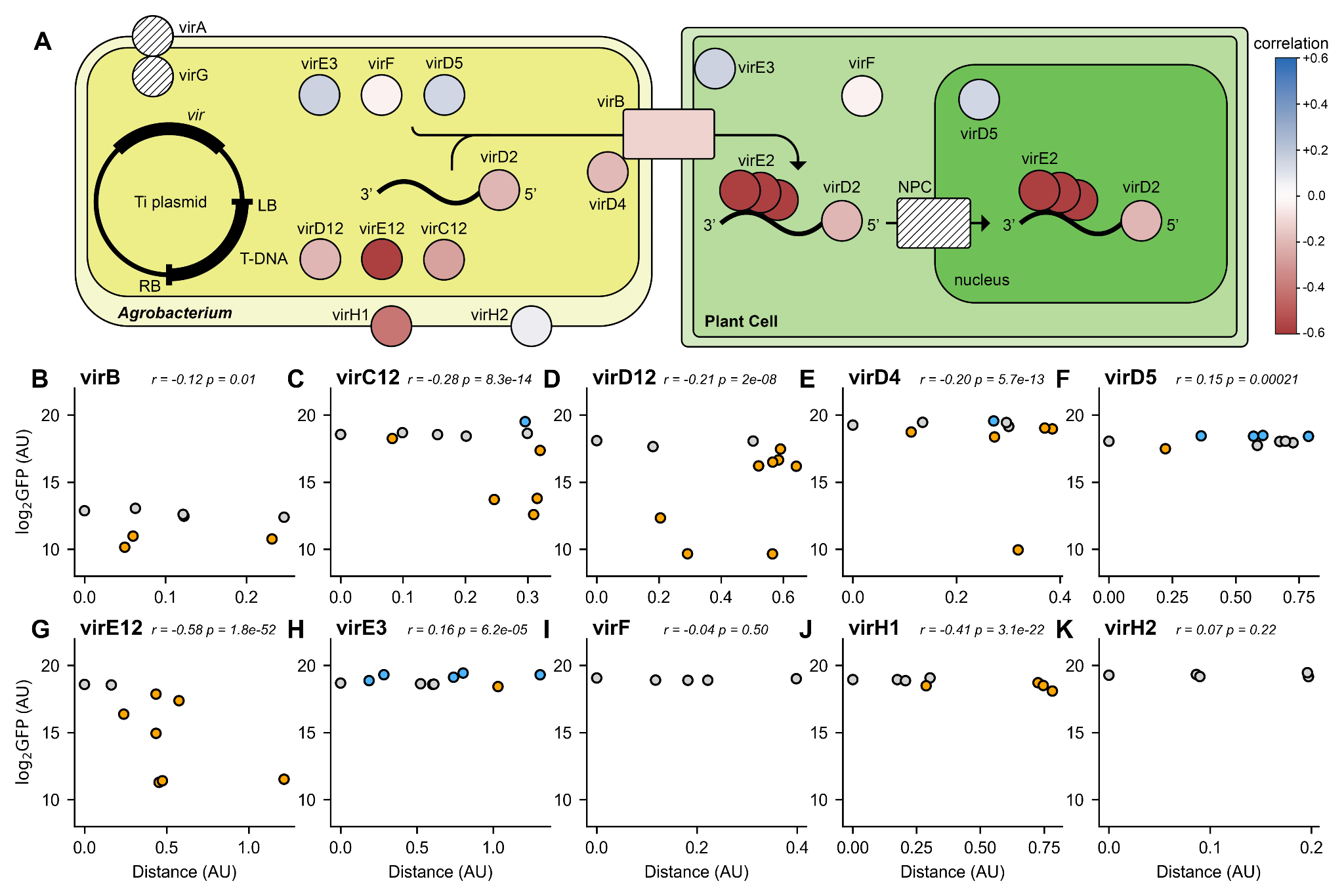

**Figure S8:** Impact of *vir* gene allele on tobacco transformation: A) Cartoon shows the localization of different *vir* gene products within the bacterial and plant cell during the AMT process. Gene products are colored by the Pearson correlation between the GFP produced in complementation assays, and the phylogenetic distance between a given allele and the wild-type *A. fabrum* C58 gene. B-K) Plots on the left show the effects of alleles of the indicated gene clusters in *A. fabrum* GV3101 deletion mutants complemented with pGinger based vectors that resulted in improved transformation. Box plots in pink show alleles that are statistically worse than the wild-type allele, box plots in blue show alleles that are statistically superior than the wild-type allele, and box plots in white show alleles that are statistically indistinguishable from the wild-type allele. Statistical significance was determined using a Bonferroni corrected T-test (p-value < 0.05, n=64). Scatterplots on the right show the results of allele complementation assays in tobacco (y-axis) as a function of phylogenetic distance of each allele from the native allele (*A. fabrum* C58). The *p*-values for both Pearson and Spearman correlation are shown above each plot.

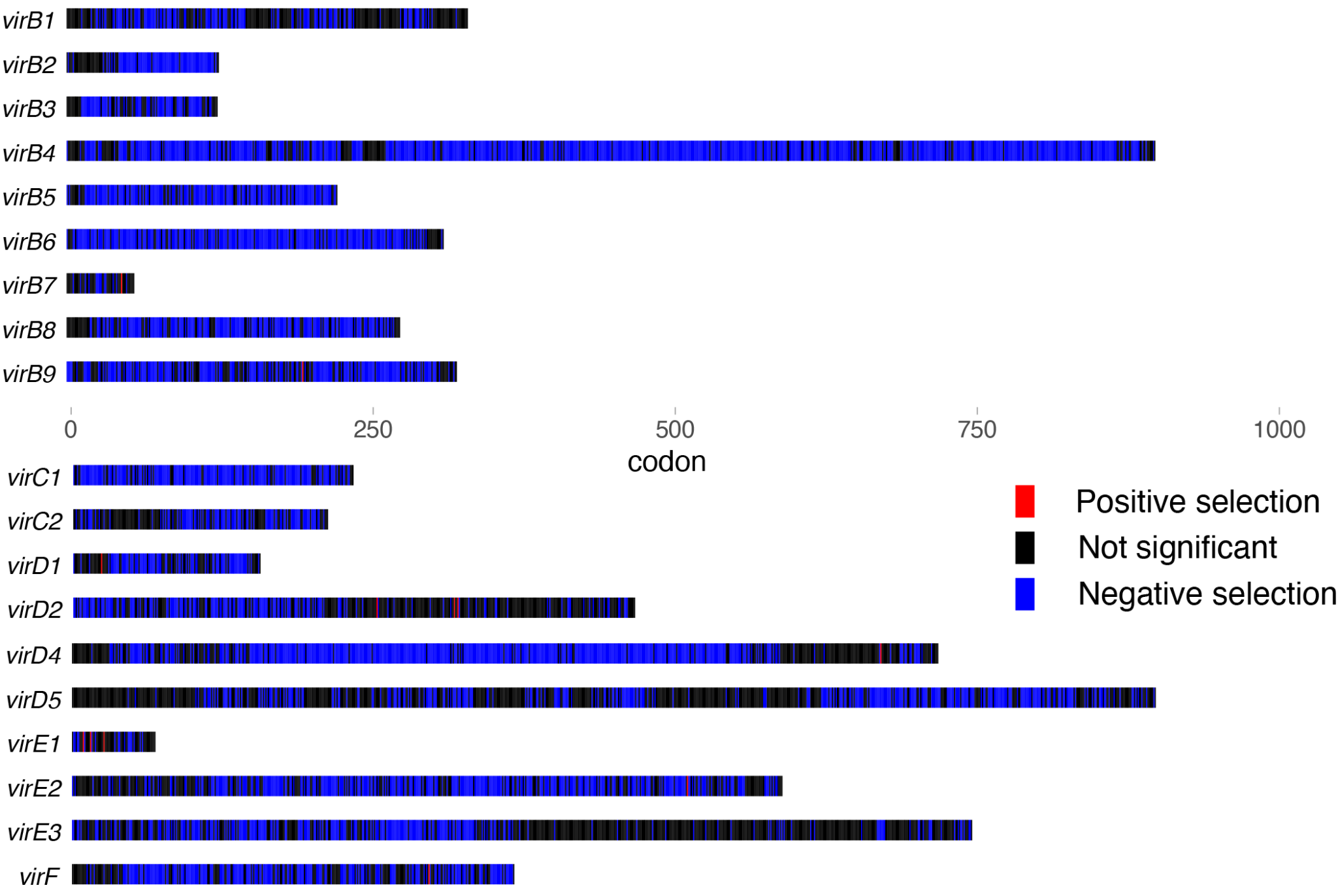

**Figure S9:** Selective pressure on *vir* genes. For each conserved *vir* gene the diversifying (positive selection shown in red) and negative (purifying selection shown in blue) selection are shown for each codon of the gene using d_N_/d_S_ analysis. Resides that have no significant selection are shown in black.

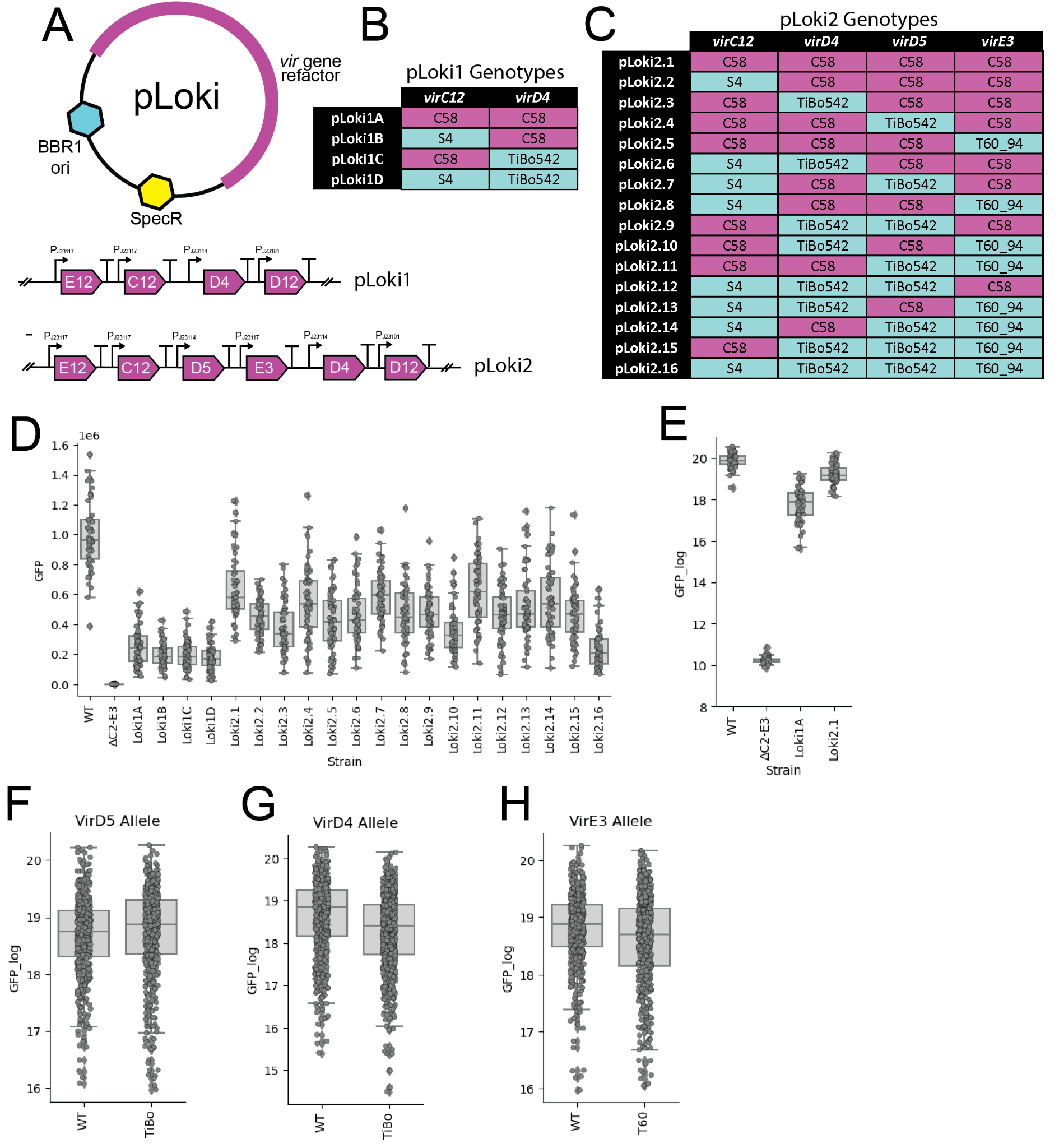

**Figure S10**: Evaluation of combinatorial *vir* allele complementation: A) Genetic Diagrams of the pLoki1.0 and pLoki2.0 vectors. B) Table shows allelic composition of pLoki1 vectors. C) Table shows allelic composition of pLoki2 vectors. D) Graph shows complementation of a *virC2*-*virE3* gene cluster deletion of *A. fabrum* GV3101 with the pLoki plasmids. The y-axis shows transient GFP production and the x-axis shows complementation with different variants (n=64). E) Graph shows complementation of a *virC2*-*virE3* gene cluster deletion of *A. fabrum* GV3101 with the pLoki plasmids harboring only wild-type (C58) alleles. The y-axis shows transient GFP production and the x-axis shows complementation with different variants (n=64). F) Composite complementation *virC2*-*virE3* gene cluster deletion with pLoki vectors that have *virD5* pTiBo542 alleles versus those harboring the wild-type *virD5* allele. G) Composite complementation *virC2*-*virE3* gene cluster deletion with pLoki vectors that have *virD4* pTiBo542 alleles versus those harboring the wild-type *virD4* allele. H) Composite complementation *virC2*-*virE3* gene cluster deletion with pLoki vectors that have *virE3* pTiT60_94 alleles versus those harboring the wild-type *virE3* allele.

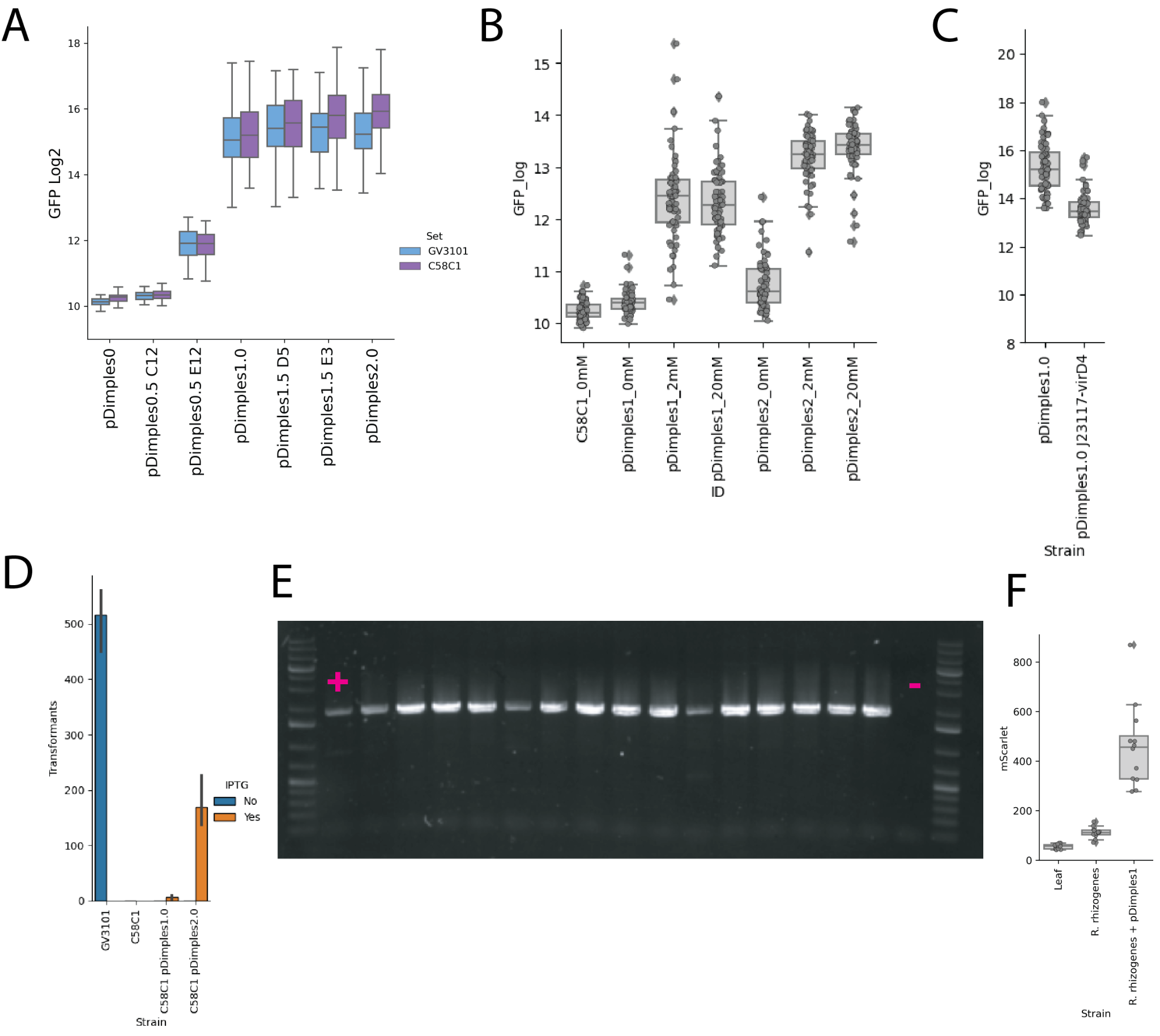

**Figure S11:** Evaluation of refactored pTi plasmids. A) Transient transformation of tobacco leaves as measured by GFP production by synthetic pTi plasmids harbored either in *A. fabrum* C58C1 or GV3101 Δ*virA-E3* (n=64). The Y-axis shows log2 transformed GFP (AU). B) Graph shows the impact of different amounts of IPTG on the ability of pDimples1.0 and pDimples 2.0 to enable tobacco transient transformation of *A. fabrum* C58C1 (n=64) C) Graph shows the impact of increasing the expression of *virD4* by replacing the PJ23114 promoter with PJ23117 in pDimples1.0 harbored by C58C1. The y-axis shows tobacco transient GFP production (n=64) D) Graph shows the number of colonies obtained by transforming *R. toruloides* with *A. fabrum* C58C1 pDimples1.0 or pDimples 2.0 with and without the addition of IPTG to induction media (n=3) E) PCR confirmation of 15 randomly selected colonies of *R. toruloides* transformed by *A. fabrum* C58C1 pDimples 2.0 with primers specific to inserted T-DNA. The first non-ladder well shows product amplified directly from binary vector plasmid, and the last non-ladder well shows negative result amplified from wild-type *R. toruloides*. F) Graph shows the amount of fluorescence from tobacco leaf punches infiltrated with either a naturally cured strain of *R. rhizogenes* or the same strain harboring pDimples1.0 (n=12).

[Bibliography](https://sciwheel.com/work/bibliography)

[11.	Song, M., Sukovich, D.J., Ciccarelli, L., Mayr, J., Fernandez-Rodriguez, J., Mirsky, E.A., Tucker, A.C., Gordon, D.B., Marlovits, T.C., and Voigt, C.A. (2017). Control of type III protein secretion using a minimal genetic system. Nat. Commun. *8*, 14737. 10.1038/ncomms14737.](https://sciwheel.com/work/bibliography/3723436)

[12.	Sorg, R.A., Gallay, C., Van Maele, L., Sirard, J.-C., and Veening, J.-W. (2020). Synthetic gene-regulatory networks in the opportunistic human pathogen Streptococcus pneumoniae. Proc Natl Acad Sci USA *117*, 27608–27619. 10.1073/pnas.1920015117.](https://sciwheel.com/work/bibliography/15366359)

[13.	Geddes, B.A., Kearsley, J.V.S., Huang, J., Zamani, M., Muhammed, Z., Sather, L., Panchal, A.K., diCenzo, G.C., and Finan, T.M. (2021). Minimal gene set from Sinorhizobium (Ensifer) meliloti pSymA required for efficient symbiosis with Medicago. Proc Natl Acad Sci USA *118*. 10.1073/pnas.2018015118.](https://sciwheel.com/work/bibliography/14871212)

[14.	Nester, E.W. (2014). Agrobacterium: nature’s genetic engineer. Front. Plant Sci. *5*, 730. 10.3389/fpls.2014.00730.](https://sciwheel.com/work/bibliography/630006)

[20.	De Saeger, J., Park, J., Chung, H.S., Hernalsteens, J.-P., Van Lijsebettens, M., Inzé, D., Van Montagu, M., and Depuydt, S. (2021). Agrobacterium strains and strain improvement: Present and outlook. Biotechnol. Adv. *53*, 107677. 10.1016/j.biotechadv.2020.107677.](https://sciwheel.com/work/bibliography/12661560)

[43.	Zhang, X., Hooykaas, M.J.G., van Heusden, G.P., and Hooykaas, P.J.J. (2022). The translocated virulence protein VirD5 causes DNA damage and mutation during Agrobacterium-mediated transformation of yeast. Sci. Adv. *8*, eadd3912. 10.1126/sciadv.add3912.](https://sciwheel.com/work/bibliography/14874743)

[44.	Zhang, X., van Heusden, G.P.H., and Hooykaas, P.J.J. (2017). Virulence protein VirD5 of Agrobacterium tumefaciens binds to kinetochores in host cells via an interaction with Spt4. Proc Natl Acad Sci USA *114*, 10238–10243. 10.1073/pnas.1706166114.](https://sciwheel.com/work/bibliography/4379255)

[45.	Wang, K., Herrera-Estrella, A., and Van Montagu, M. (1990). Overexpression of virD1 and virD2 genes in Agrobacterium tumefaciens enhances T-complex formation and plant transformation. J. Bacteriol. *172*, 4432–4440. 10.1128/jb.172.8.4432-4440.1990.](https://sciwheel.com/work/bibliography/1340720)

[46.	Weisberg, A.J., Davis, E.W., Tabima, J., Belcher, M.S., Miller, M., Kuo, C.-H., Loper, J.E., Grünwald, N.J., Putnam, M.L., and Chang, J.H. (2020). Unexpected conservation and global transmission of agrobacterial virulence plasmids. Science *368*. 10.1126/science.aba5256.](https://sciwheel.com/work/bibliography/9127618)

[47.	Weisberg, A.J., Wu, Y., Chang, J.H., Lai, E.M., and Kuo, C.H. (2023). Virulence and Ecology of Agrobacteria in the Context of Evolutionary Genomics. Annu. Rev. Phytopathol. *61*.](https://sciwheel.com/work/bibliography/14434235)

[55.	Ham, T.S., Dmytriv, Z., Plahar, H., Chen, J., Hillson, N.J., and Keasling, J.D. (2012). Design, implementation and practice of JBEI-ICE: an open source biological part registry platform and tools. Nucleic Acids Res. *40*, e141. 10.1093/nar/gks531.](https://sciwheel.com/work/bibliography/1192584)

[56.	Chen, J., Densmore, D., Ham, T.S., Keasling, J.D., and Hillson, N.J. (2012). DeviceEditor visual biological CAD canvas. J. Biol. Eng. *6*, 1. 10.1186/1754-1611-6-1.](https://sciwheel.com/work/bibliography/1030617)

[57.	Hillson, N.J., Rosengarten, R.D., and Keasling, J.D. (2012). j5 DNA assembly design automation software. ACS Synth. Biol. *1*, 14–21. 10.1021/sb2000116.](https://sciwheel.com/work/bibliography/312499)

[58.	Gibson, D.G., Young, L., Chuang, R.-Y., Venter, J.C., Hutchison, C.A., and Smith, H.O. (2009). Enzymatic assembly of DNA molecules up to several hundred kilobases. Nat. Methods *6*, 343–345. 10.1038/nmeth.1318.](https://sciwheel.com/work/bibliography/30022)

[59.	Engler, C., Kandzia, R., and Marillonnet, S. (2008). A one pot, one step, precision cloning method with high throughput capability. PLoS ONE *3*, e3647. 10.1371/journal.pone.0003647.](https://sciwheel.com/work/bibliography/706971)

[60.	Green, M.R., and Sambrook, J. (2012). Molecular Cloning: A Laboratory Manual (Fourth Edition), Volume 1, 2 & 3 4th ed. (Cold Spring Harbor Laboratory Press).](https://sciwheel.com/work/bibliography/15144211)

[66.	Gin, J., Chen, Y., and J Petzold, C. (2020). Chloroform-Methanol Protein Extraction for Gram-negative Bacteria (High Throughput) v1. 10.17504/protocols.io.bfx6jpre.](https://sciwheel.com/work/bibliography/15144207)

[67.	Chen, Y., Gin, J., and J Petzold, C. (2021). Discovery proteomic (DDA) LC-MS/MS data acquisition and analysis v2. 10.17504/protocols.io.buthnwj6.](https://sciwheel.com/work/bibliography/13431560)

[74.	Jones, E., Oliphant, T., Peterson, P., and Others SciPy: Open source scientific tools for Python.](https://sciwheel.com/work/bibliography/5778430)

[75.	Gramfort, A., Luessi, M., Larson, E., Engemann, D.A., Strohmeier, D., Brodbeck, C., Parkkonen, L., and Hämäläinen, M.S. (2014). MNE software for processing MEG and EEG data. Neuroimage *86*, 446–460. 10.1016/j.neuroimage.2013.10.027.](https://sciwheel.com/work/bibliography/1040961)

[76.	Katoh, K., and Standley, D.M. (2013). MAFFT multiple sequence alignment software version 7: improvements in performance and usability. Mol. Biol. Evol. *30*, 772–780. 10.1093/molbev/mst010.](https://sciwheel.com/work/bibliography/387873)

[77.	Price, M.N., Dehal, P.S., and Arkin, A.P. (2010). FastTree 2 — approximately maximum-likelihood trees for large alignments. PLoS ONE *5*, e9490. 10.1371/journal.pone.0009490.](https://sciwheel.com/work/bibliography/178753)

[78.	Sukumaran, J., and Holder, M.T. (2010). DendroPy: a Python library for phylogenetic computing. Bioinformatics *26*, 1569–1571. 10.1093/bioinformatics/btq228.](https://sciwheel.com/work/bibliography/396911)
